# CATHe: Detection of remote homologues for CATH superfamilies using embeddings from protein language models

**DOI:** 10.1101/2022.03.10.483805

**Authors:** Vamsi Nallapareddy, Nicola Bordin, Ian Sillitoe, Michael Heinzinger, Maria Littmann, Vaishali Waman, Neeladri Sen, Burkhard Rost, Christine Orengo

## Abstract

CATH is a protein domain classification resource that combines an automated workflow of structure and sequence comparison alongside expert manual curation to construct a hierarchical classification of evolutionary and structural relationships. The aim of this study was to develop algorithms for detecting remote homologues that might be missed by state-of-the-art HMM-based approaches. The proposed algorithm for this task (CATHe) combines a neural network with sequence representations obtained from protein language models. The employed dataset consisted of remote homologues that had less than 20% sequence identity. The CATHe models trained on 1773 largest, and 50 largest CATH superfamilies had an accuracy of 85.6+−0.4, and 98.15+−0.30 respectively. To examine whether CATHe was able to detect more remote homologues than HMM-based approaches, we employed a dataset consisting of protein regions that had annotations in Pfam, but not in CATH. For this experiment, we used highly reliable CATHe predictions (expected error rate <0.5%), which provided CATH annotations for 4.62 million Pfam domains. For a subset of these domains from homo sapiens, we structurally validated 90.86% of the predictions by comparing their corresponding AlphaFold structures with experimental structures from the CATHe predicted superfamilies.

## 2. Introduction

The CATH database (www.cathdb.info) (1) classifies protein domain structures into superfamilies when there is strong evidence that domains share a common evolutionary ancestor. The resource was established in 1994 and currently comprises over 500,000 domains from experimental structures in the Worldwide Protein Data Bank (wwPDB)(2), classified into 5,481 CATH superfamilies. This information has been used to predict domain locations and superfamilies in over 150 million protein sequences (3). The first step in the classification process is to chop the 3D structures deposited to the wwPDB into their constituent structural domains, using domain detection algorithms that either detect structural similarity with classified domains or detect compactly folded regions in the chain (1). Extensive manual curation is used to validate domain boundaries for distant homologues and novel folds. The classification protocol exploits a number of computational methods to provide evidence for evolutionary relationships including structure-based (SSAP (4), and CATHEDRAL (5)), and sequence-based comparison tools (HMMER (6), PRC (7), HHsearch (8), and MMseqs2 (9)).

3D protein structure information is often crucial for understanding protein function and functional mechanisms, and can also help to identify extremely remote evolutionary relationships (10). Since experimental structural data is only available for a small (< 1%) fraction of known protein sequences, it can be useful to employ sequence-based tools to detect CATH homologues from large protein sequence resources, such as UniProt (11) and ENSEMBL (12). This expands the available sequence, functional and evolutionary data in each superfamily. Furthermore, the recent DeepMind announcement stating that they are going to provide AlphaFold2 (13) predictions for the whole of UniRef90 will provide an excellent opportunity to classify more domains into CATH using structure comparison tools. Structure comparison methods for detecting remote homologues tend to be much slower than sequence-based methods and therefore improved sequence strategies for remote homologue detection could reduce the time for assigning these models to the appropriate superfamily.

CATH homologous superfamilies are organized into a hierarchy of Class, Architecture, and Topology reflecting similarities in secondary structure composition, the orientation of secondary structures, and folding, respectively. CATH provides two levels of release: CATH-B, which is a daily snapshot of the latest structural domain boundaries and superfamily assignments, and CATH+, which adds layers of derived data, such as the predicted sequence domains, functional annotations, and functional clustering. The latest version of CATH+ (version 4.3) classifies 151 million predicted sequence domains into 5,481 superfamilies. Additional functional information from UniProtKB (data such as their Gene Ontology (GO) (14) annotations), Catalytic Site Atlas (15), along with Enzyme Commission (16) numbers are provided.

CATH is widely accessed by the biology and computational biology community, and the classification of domains into evolutionary superfamilies has been valuable in revealing the unequal population of superfamilies whereby the top 200 superfamilies account for about 64% of the non-redundant CATH domains (17). It has also enabled detailed studies of structural mechanisms of functional divergence in proteins during evolution (18) and revealed sites under positive selection implicated in antimicrobial resistance (19). Recent studies exploited functional families in CATH to detect sites involved in the binding of SARS-CoV-2 Spike protein to the ACE2 receptor in humans and other animals (20).

In some superfamilies, relatives can diverge significantly with residue insertions leading to a 3-fold variation in size, or more, making it hard to align them well and detect their evolutionary relationship (18). Although HMM-HMM strategies like HHblits (21) have proven powerful in detecting very remote homologues, even where significant structural divergence exists, methods that improve on HHblits in sensitivity and speed would allow CATH and other related resources to keep pace with the expansion in the protein sequence repositories, including MGnify (22), and the Big Fantastic Database (BFD) (23, 24) which are about 20 fold, and 11 fold larger than UniProt respectively.

We recently explored the value of methods using vector representations (embeddings) of protein sequences obtained from the protein language model (pLM) ProtBERT (25) to detect functional similarity between proteins (26). In this study, we employed an embedding strategy more powerful than ProtBERT to capture sequence relationships within CATH superfamilies and exploit these to recognize even very remote homologues (i.e., less than 20% sequence identity). Our new approach (CATHe; short for CATH embedding) uses protein sequence embeddings from state-of-the-art language models (LMs) adapted from natural language processing (NLP). Instead of being trained on natural language, pLMs were trained on large protein sequence sets from UniProt. The so-called self-supervised pre-training (a special form of unsupervised training) of those pLMs allows them to readily derive benefits from large unlabeled data as they are only trained on reconstructing corrupted input tokens (words/amino acids) from non-corrupted sequence context (sentence/protein sequence). In a second step, the learned information can be transferred to downstream tasks (transfer learning) such as prediction of protein structure (27), function (28), and prediction of single amino acid variant effect (29). As pLMs see no other labels during pre-training but masked amino acids, there is no risk of information leakage between the first stage pre-training and second stage supervised training. In this work, embeddings from one pLM (ProtT5(25)) were used to train machine learning models to classify protein sequences into CATH superfamilies. To make sure that the developed models were capable of detecting very remote homologues in these superfamilies, the models were built to detect non-redundant homologues with at most 20% sequence identity.

Our work builds on earlier studies that have exploited machine learning approaches to classify protein sequences into families. The ProtENN (30) method used an ensemble deep learning framework that generates protein sequence embeddings to classify protein domains into Pfam (31) families. The ensemble was built from 13 variations of the ProtCNN models which were developed using the ResNet (32) architecture. ProtENN achieved an accuracy of 87.8% when the sequence identity between testing and training sets was set to be less than 25%. Bepler and Berger (33) used sequence information extracted from pLMs in conjunction with structural information to predict SCOP families derived from the ASTRAL benchmark dataset (34) which uses a sequence redundancy filter of 40%. Their proposed model, the MT-LSTM, achieved an accuracy of 96.19%. Additionally, they found that including structural data led to a much better organization of the proteins in the embedding space when compared to the use of sequence information alone. An important conclusion from this study was the fact that pLMs had the power to capture structure, function, and evolutionary information from sequence alone.

DeepFam (35) is another such deep learning based study on classifying protein sequences from the Clusters of Orthologous Groups of proteins (COG) database and the G Protein-Coupled Receptor (GPCR) dataset. DeepFam did not employ a pLM to generate protein sequence embeddings but instead used one-hot encodings of the sequence as input for a neural network model, furthermore, there was no application of a sequence redundancy filter on their dataset. DeepFam attained a prediction accuracy of 97.17% on the family level on the GPCR dataset, and 95.4% on the COG dataset (with protein families having at least 500 sequences). DeepNOG (36) employed a method similar to DeepFam to classify sequences from COG and eggNOG5 databases. Furthermore, deep learning techniques have been used not just for family-level classification, but also for fold recognition, in FoldHSphere (37). In this study, deep residual networks were employed to generate protein sequence embeddings optimized for fold recognition in SCOP.

To our knowledge, we have used a more stringent test set than these approaches to validate CATHe, where we used a redundancy filter of 20% sequence identity. Our model was able to predict the correct CATH superfamily with an F1-Score of 72.3% +−0.6%, and an Accuracy of 85.6+−0.4. In an effort to examine the potential of CATHe when it comes to detecting remote homologues better than the current HMM (Hidden Markov Models) approaches, we analyzed regions of protein sequences that had been classified by Pfam but not CATH (i.e. sequence domains that were not detected by the existing CATH HMM library). Using only predictions up to an expected error rate of 0.5%, CATHe was able to provide CATH superfamily annotations for an additional 4.62 million Pfam domains. Using a subset of these Pfam domains in *homo sapiens* that had good quality AlphaFold models, we were able to structurally validate 90.86% of the CATHe predictions using SSAP (4) (short for Sequential Structure Alignment Program, which is a tool for structure comparison).

## 3. Results

For the process of developing a machine learning model, we generated a dataset using the CATH (version 4.3) database. The training set had domains from the whole database (i.e., CATH relatives with known structure from PDB and those with predicted CATH structural annotations, from UniProt), whereas the testing and validation sets consisted only of domains of known structure in CATH. To be certain that the models learned to detect remote homologues, we made sure that there was less than 20% sequence identity between and within the three sets. In order to convert the domains into a format accessible for the machine learning models, we generated sequence embeddings for all of them. We tested the performance of two machine learning models (Artificial Neural Networks (ANNs), logistic regression) trained on embeddings from the ProtBERT and ProtT5 pLMs and simple strategies designed as a control (e.g., homology based inference based on BLAST, prediction based on protein sequence length, and a random baseline). Please refer to the Materials and Methods section for a more detailed description of the models.

### 3.1 Evaluation on the TOP 1773 SUPERFAMILIES dataset

The TOP 1773 SUPERFAMILIES dataset consisted of 1773 CATH superfamilies (comprising of superfamilies with at least 2 non-redundant relatives in the PDB) and the Other superfamilies set (comprising of 3456 superfamilies having less than 2 PDB relatives in the superfamily). This was chosen as a negative control to ensure that the model prediction probabilities were not skewed towards more highly populated superfamilies. The superfamilies from the TOP 1773 SUPERFAMILIES dataset comprise nearly 90% of all non–redundant CATH domains (see Methods for details). This dataset was randomly split into three subsets, for training, validation, and testing, ensuring that all proteins had a maximum of 20% sequence identity within and between each set (for more details on the generation of the TOP 1773 SUPERFAMILIES dataset, please refer to section 5.1.). These sets were used to explore the performance of seven prediction models: 1.) homology-based inference (HBI) via BLAST, 2) an ANN trained on ProtBERT embeddings, 3.) CATHe (ANN model trained on sequence embeddings from ProtT5), 4.) an ANN trained only on sequence lengths, 5.) logistic regression (LR) trained on ProtBERT embeddings, 6.) LR trained on ProtT5 embeddings, and 7.) a random baseline (for details on the development of the seven prediction models, please refer to section 5.4, Table S7). The performance of the seven prediction models was measured on the testing set of the TOP 1773 SUPERFAMILIES dataset using accuracy, F1-Score, MCC, and Balanced Accuracy (for more information on the metrics, please refer to section S2.3). The results have been provided in detail in the supplementary (section S1.1.).

When comparing the seven methods, the two models trained on ProtT5 embeddings (CATHe and LR(ProtT5)) outperformed all other methods, including those trained on ProtBERT (ANN (ProtBERT) and LR (ProtBERT)). Among models trained on ProtT5, the ANN outperformed the LR with an F1-Score of 72.3% +−0.6%. Compared to these embedding-based approaches, HBI via BLAST reached lower performance but outperformed the ANN trained on sequence lengths which improved only slightly over the random baseline. In general, we can see that the models trained on embeddings from pLMs outperformed traditional methods like BLAST.

### 3.2. Evaluation on the largest 50 superfamilies

The TOP50 SUPERFAMILIES dataset (please refer to section 5.1.2 for details on how this dataset was generated) is a subset of the TOP 1773 SUPERFAMILIES dataset, consisting only of the 50 largest superfamilies chosen according to the population of the superfamily in the TOP 1773 SUPERFAMILIES dataset. These are highly populated superfamilies accounting for 37.32% of all non-redundant domains in CATH. We re-trained seven different models (refer to Table S7) and measured their performance on the testing set for the TOP50 SUPERFAMILIES dataset using four metrics, Accuracy, F1-Score, MCC, and Balanced Accuracy. The results are illustrated in Fig 1 (B) and described in detail in Table S1 (B).

**Fig 1.**
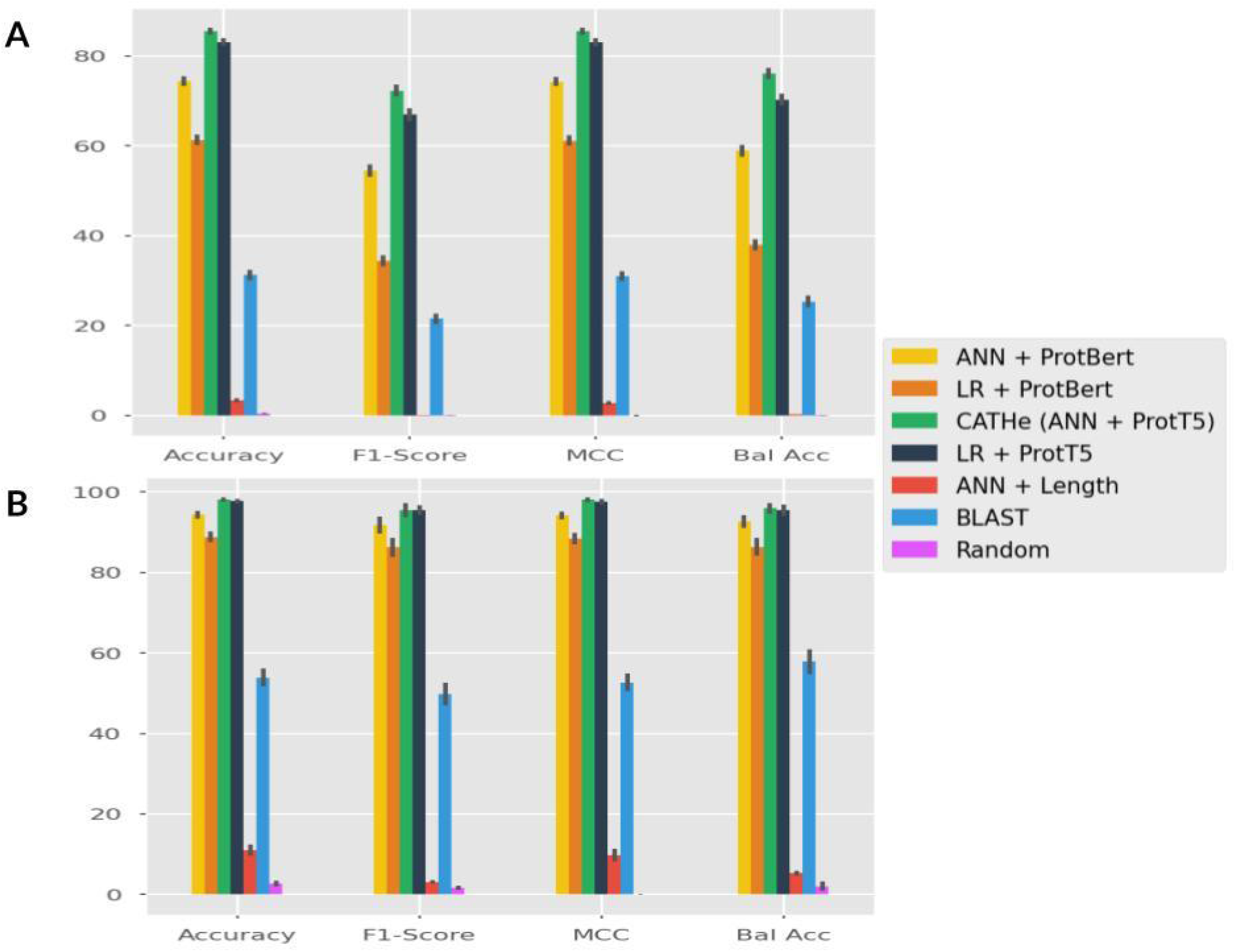
(A) Performance comparison using TOP 1773 SUPERFAMILIES. Seven methods for remote homology detection for CATH homologous superfamilies were compared using the TOP 1773 SUPERFAMILIES test set: two models trained on ProtT5 embeddings (ANN and LR with the ANN being dubbed CATHe), two models trained on ProtBERT embeddings (ANN and LR) and three baselines consisting of homology-based inference via BLAST, an ANN trained only on protein sequence lengths and a random baseline. The performance was measured using Accuracy, F1-Score, MCC, and Balanced Accuracy (Bal Acc) along with 95% confidence intervals. The proposed model, CATHe, has the highest performance, and the machine learning models that were trained on pLM embeddings outperformed BLAST. (B) Performance comparison using TOP 50 SUPERFAMILIES dataset, the previously mentioned approach with the seven methods and four metrics were used in this analysis as well. We notice a similar trend that the proposed CATHe model had the highest performance, and the machine learning models trained on pLM embeddings had a better performance than the baselines (including BLAST).

Again, the highest performance (F1-Score of 95.4% +−0.9%) among all these seven models was achieved by CATHe. The performance of the other six models follows a similar trend as they did in the results of the TOP 1773 SUPERFAMILIES dataset, i.e., models trained on ProtT5 outperform models trained on ProtBERT and pLM-based methods outperform baselines, including HBI via BLAST.

To understand how superfamilies are organized in embedding space, and to which extent CATHe picked up superfamily-specific information during training, we used t-SNE (38) to visualize high-dimensional embedding space in 2-D. Towards this end, t-SNE was used to project the 1024-D ProtT5 embeddings for all protein domains in the TOP 50 SUPERFAMILIES test set down to 2-dimensional space. For the same set, 128-D CATHe embeddings were derived from the final layer of the CATHe ANN after training and also projected down to 2-D using t-SNE.

Coloring the 2-D projections (Fig. 2A) based on CATH architecture highlights that embeddings from the ProtT5 pLM are able to cluster some of the architectures such as 2.60, 2.40, and 1.10 quite well, but the larger, more diverse architectures such as 3.40 and 3.30 are more dispersed. One possible explanation for this is that there are many different topologies for the superfamilies in these latter two architectures (126 for 3.40 and 224 for 3.30, compared to <50 for most of the remaining architectures). Furthermore, these latter two architectures comprise superfamilies that are particularly structurally diverse, in some cases showing 3-fold or more variation in the size of the relatives. In figures S4 and S5 we compare two domains from the architectures 3.30 and 3.40 respectively to show the amount of structural diversity in these architectures. The denser clustering of CATH architectures in CATHe embedding space (Fig. 2 (B)) indicates that CATHe is able to pick up on differentiating information on the architecture level during the training process. CATHe is also capable of picking up on fold-level information, we can see in figure S6 that the point clouds for the different topologies in the 3.40 architecture are well separated.

**Fig 2.**
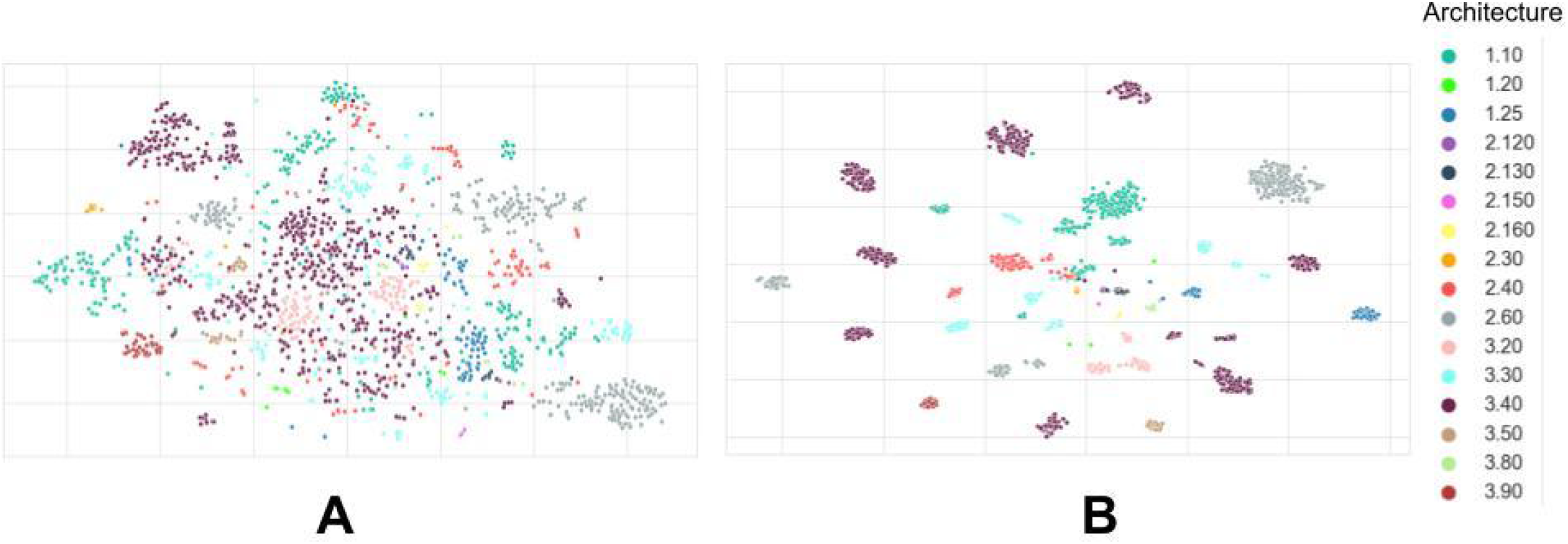
T-SNE projections of high-dimensional embedding spaces for the TOP50 SUPERFAMILIES test set colored by CATH architectures. Panel (A) shows the information learned by the pLM ProtT5 during self-supervised pre-training, i.e., without any supervised training. For panel (B), embeddings were extracted from the last layer of the trained CATHe ANN to highlight the additional benefit of supervised training for distinguishing CATH architectures.

### 3.3. Class imbalance studies

A number of tests were conducted to understand how the class imbalance for homologous superfamilies in the two datasets, TOP 1773 SUPERFAMILIES, and the TOP50 SUPERFAMILIES, affected the CATHe performance. The results for those tests are shown in Figures S1 and S2 for the TOP 1773 SUPERFAMILIES dataset and the TOP50 SUPERFAMILIES dataset, respectively.

Examining the relationship between the population of a superfamily as defined by the number of sequences for this superfamily in the training set and the CATHe performance for these superfamilies in terms of F1-Score (Fig. S1A-C, S2A-C) showed that no linear relationship exists between the two. Further analysis of the relationship between structural diversity of relatives as defined by the number of Structurally Similar Groups at 5 Å (SSG5 groups, see S1.1. for more details) and CATHe performance (Fig. S1D, S2D) also does not suggest a direct linear relationship. For more information see sections S1.1., and S1.2. in the supplementary.

### 3.4. Applications of CATHe

The dataset used for analyzing CATHe consisted of domains that are present in Pfam v 33.1 but could not be transferred into CATH v 4.3 using the current Hidden Markov Model (HMM) approaches for CATH superfamily classification. In the approach to generate this list of domains, the first step was to scan protein sequences from UniProt against the HMM libraries derived from CATH superfamilies (generated by running HMMER3 on all non-redundant representative sequences at 95% sequence identity from the CATH superfamilies). The regions from the first step which did not match with the CATH HMMs were scanned against the library of HMMs derived from Pfam (v 33.1), and the ones that matched with the Pfam HMMs were used to build the Pfam domains list. The CATHe model trained on the TOP 1773 SUPERFAMILIES dataset was used to analyze this Pfam domains list. Additionally, we conducted an in-depth analysis on a subset of these Pfam domains which belonged to *homo sapiens*.

#### Total Pfam Domains

There are nearly 38M domains present in Pfam which could not be detected by CATH HMMs. These domains were clustered at 60% sequence identity using MMseqs2. This threshold was chosen as it has been shown to cluster relatives with significant structural and functional similarities (39). The resulting *PFAM-S60* set had 10.5M domains which were converted to embeddings using ProtT5. Before predicting using CATHe, its error rate was monitored through a threshold analysis using CATHe predictions on the validation set of the TOP 1773 SUPERFAMILIES dataset. From these predictions, we measured the error rate at various prediction probability thresholds (see S1.1 for more details). From these analyses, we concluded that only considering predictions with a prediction probability above 0.9 (or 90%, which corresponds to a 49.75% coverage) allows us to keep the expected error rate at 0.5%,

Applying this threshold to CATHe predictions for the PFAM-S60 set allowed us to assign CATH homologous superfamilies to, roughly 4.6M (4,623,730) PFAM-S60 domains (43.72% of the PFAM-S60 set). Adding these newly predicted domains to CATH v 4.3 led to an increase in database size of 3.06%.

#### Human Pfam Domains

We considered comparing CATHe against state-of-the-art HMM-based techniques. However, conducting a direct comparison between the performance of these two techniques on the same dataset would mean building HMM libraries from the sequences of training sets of the TOP 1773 SUPERFAMILIES and TOP 50 SUPERFAMILIES datasets. This would result in extremely poor performing HMMs as the alignments from which the HMMs would be derived would be built from a small set of sequences that had <20% sequence identity (refer to sections 5.1 and 5.2 for more information on how the datasets were developed). These very small training sets make it challenging to perform a direct and fair comparison. Therefore, we chose to compare CATHe and HMM approaches by taking the CATHe predictions, for domains not matched by HMMs, and structurally validating them against known structures in the CATHe predicted CATH superfamily.

Therefore, we chose a subset of Pfam domains (not detected by CATH-HMMs) belonging to *homo sapiens* (list of 36k domains). We extracted the appropriate domain structures from the AlphaFold2 (AF2) models obtained from the EBI database (https://alphafold.ebi.ac.uk/) (13, 40). From this set, we removed those domains that had problematic regions or were predicted to be low-quality models (refer to Section 5.3. for more information). Choosing those domains for which CATHe gave a good prediction probability (probability threshold of 90%; 0.5% error rate), resulted in 197 domains (dubbed *PFAM-human*).

To structurally validate the CATHe superfamily predictions for PFAM-human, we used our in-house protein structure comparison method, SSAP (4). The Pfam-human regions were extracted from AF2 predicted structures, then scanned against the structural domains from the CATHe predicted superfamily. However, since these are likely to be remote homologues of CATH superfamilies, the CATH superfamilies were first expanded (by ~60%, on average) by including AF2 predicted structural regions corresponding to close CATH-HMM matches. These CATH-HMM matches were validated by structure comparison (using a strict SSAP score threshold of 80 and a domain overlap threshold of 60%) (see Methods for more details). The AlphaFold structures of PFAM-human domains were then scanned against the expanded CATH superfamilies using SSAP.

To identify acceptable structural matches for classification in a CATH superfamily, we applied individual SSAP score thresholds for the different CATH classes predicted by CATHe (SSAP score thresholds of 71, 66, and 69 for CATH classes 1, 2, and 3 respectively). This SSAP score threshold was applied in addition to a structural overlap threshold of 60. These thresholds are associated with a 5% error rate (refer to Supplementary section S2.4. for details on how we arrived at these thresholds). Applying these thresholds, we are able to bring 142 domains into CATH from the 197 domains in Pfam-human. Additionally, we conducted manual curation on the 55 domains that didn’t cross the thresholds and noticed that 37 more domains match at the superfamily level (refer to supplementary S1.1. for more information on the manual curation). Therefore a total of 179 domains out of 197 (90.86%) from Pfam-human match at the superfamily level and can be safely brought into CATH.

In order to examine differences in AF2 domain predictions made by CATHe and CATH-HMM, we compared the results when using SSAP to search these predicted domains against their assigned superfamilies (fig 3). For both CATHe and CATH-HMM, we only considered comparisons involving *homo sapiens* AF2 models where the pairwise overlap from SSAP was greater than 60%. For CATH-HMMs, we only included comparisons involving domains predictions with high-quality AF2 structures (see section 5.1.3). For CATHe, this filtering left 150 domains (from 197); for CATH-HMMs this left 2,069 domains (from 39,405). The same conditions and thresholds were applied to both sets so that we could compare both of them at the same level. We observed that the CATH-HMM set of domains predicted to belong to CATH superfamilies had a greater average SSAP score (89.45) than the CATHe set (84.60) (fig. 3), which is understandable as the CATH-HMMs bring in closer homologues. CATHe was trained on the CATH-HMM predicted domains and tries to go beyond the set of closer homologues to detect remote homologues, hence more dispersed SSAP scores are expected.

**Fig 3.**
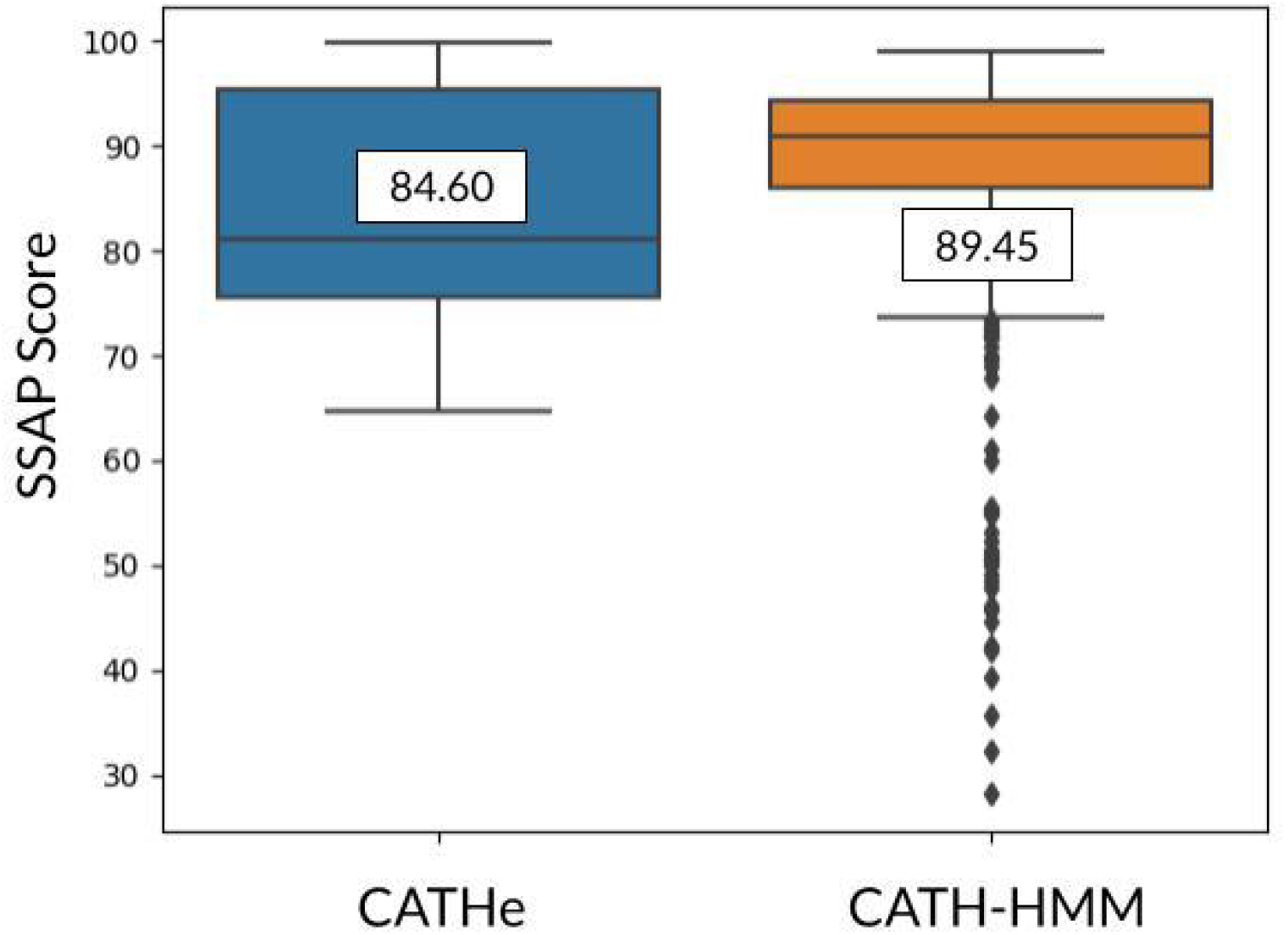
Comparison of SSAP scores for superfamily predictions made by the CATH-HMMs for Gene3D domains from *homo sapiens* (orange box) and CATHe for Pfam domains from *homo sapiens* (blue box). The SSAP boxplots for both of these sets have been annotated with their average SSAP score. The CATH-HMM set had a greater average SSAP score than the Pfam-human set showing that they tend to be closer homologues to the existing CATH superfamily relatives than those detected by CATHe.

We selected one of the CATHe matches to assess the value of these new assignments, the “Q8NE79_40_267” domain (Q8NE79: Blood vessel epicardial substance protein (residue range 40 to 267). This domain was assigned to the CATH superfamily 2.60.120.10 by CATHe with a 100% prediction probability and was subsequently manually curated to confirm the superfamily match. This domain was aligned with the sequences in the CATHe assigned superfamily, using jackhmmer (41) with an e-value cutoff of 1e-5, set for 3 iterations. The scorecons (42) tool was used to identify the conserved residues from the alignment. Residues 165, 179, and 200 were found to be conserved (scorecons score > 0.8). Furthermore, we observed that there is a mutation in residue 201, Ser201Phe, which lies near the highly conserved residues, which leads to limb-girdle muscular dystrophy autosomal recessive 25 (LGMDR25). Serine 201 was also noticed to be well conserved with a scorecons score of 0.786.

## 4. Discussion and Conclusion

The CATH database currently uses a combination of protein structure comparison tools (SSAP, CATHEDRAL) and sequence comparison tools (BLAST, HMMER, PRC, HHsearch, and MMseqs2) to identify homologues for CATH superfamilies. The predicted domains present in the CATH superfamilies were obtained using HMM-based approaches. In this study, we present a deep learning based tool, CATHe, that trains an artificial neural network (ANN) on embeddings from the protein Language Model (pLM) ProtT5 to detect remote homologues for these CATH superfamilies.

CATHe was trained, optimized, and evaluated on three datasets (training, validation, testing) generated from the CATH database and with less than 20% sequence identity and 60% overlap, within the sets and between them. The CATHe model attained an accuracy of 85.6 +− 0.4, and an F1-Score of 72.3% +− 0.6% on predicting the most fine-grained level (homologous superfamily) from the TOP 1773 SUPERFAMILIES dataset (this dataset has 1773 of the largest CATH superfamilies in addition to another set which consists of smaller superfamilies, termed as “Other” superfamilies) (Table S1 (A)). In addition, with the TOP50 SUPERFAMILIES (which has the largest 50 superfamilies from the TOP 1773 SUPERFAMILIES dataset), it was able to achieve an even higher accuracy of 98.1% +− 0.3% (Table S1 (B)). Further analysis of the CATHe model revealed that the performance of the model on an individual superfamily was not in a linear relationship with the size of the superfamily nor its structural diversity (Fig S1 and S2).

### Datasets

We developed two CATHe models for classifying protein domain sequences into CATH superfamilies (refer to section 5.2 for more information on the classification models used). These were built from two datasets, TOP 1773 SUPERFAMILIES, and TOP 50 SUPERFAMILIES. The datasets we generated to train CATHe are very stringent, as we applied multiple levels of redundancy removal to make sure that we have a dataset that properly encapsulates the features of remote homologues (refer to Section 5.1. for details on data processing). In particular, we used a very low threshold on redundancy (<= 20% sequence identity). This is a much stricter threshold than applied in related studies on protein sequence classification such as DeepFam (35) and DeepNOG (36) which did not apply any redundancy removal filters, and Bepler and Berger’s study (33) that used a sequence identity filter of 40%, but CATHe achieved a comparable performance.

### Trends in the results

The results on TOP 1773 SUPERFAMILIES and the TOP50 SUPERFAMILIES follow a similar trend. The CATHe model has the best performance, and the Random model has the worst performance. The models trained on ProtT5 embeddings (ANN dubbed CATHe, and Logistic Regression (LR)) outperformed the ones trained on ProtBERT embeddings (ANN, and LR). The results on the TOP 1773 SUPERFAMILIES dataset show that the ANN models (CATHe, and ANN(ProtBERT)) perform significantly better than the two LR models (LR(ProtT5), and LR(ProtBERT)), justifying the usage of an ANN which has a higher number of parameters as compared to a logistic regression model.

Furthermore, from the low performance of the ANN model trained on sequence lengths (ANN + Sequence Length), we conclude that sequence length is not a feature that is very useful for superfamily classification. The significant improvement in the performance of the ANN models trained on sequence embeddings (CATHe, and ANN + ProtBERT) over ANN + Sequence Length suggests that these models not only capture sequence features such as length but are also able to extract superfamily specific information from the sequence.

### In-depth analysis of the results on the TOP 50 SUPERFAMILIES dataset

The CATHe model made a total of 36 misclassifications on the testing set of the TOP50 SUPERFAMILIES dataset (Table S2). We analyzed the misclassifications with respect to the four levels of the CATH hierarchy. Even when the superfamily annotation was incorrect, CATHe was able to correctly predict the Class (level one of CATH hierarchy) and Architecture (level two of CATH hierarchy) for 23 and 14 domains respectively (refer to supplementary S1.2 for more information). This suggests that CATHe is able to capture elements of secondary structures and the 3D packing of secondary structures.

### Application of CATHe to Pfam and structural validation of the predictions

In order to compare the performance of CATHe to HMM-based sequence classification methods, we applied CATHe to a set of Pfam domains that effectively correspond to regions of UniProt sequences not recognized by the existing CATH HMM protocols (with an e-value threshold of 1e-3). For those which CATHe assigned to a CATH superfamily and which had a good quality 3D model generated by AlphaFold, with no problematic features (see section 5.3. for more details), we validated the match by performing structural comparisons of the Pfam domain’s AlphaFold model against relatives from the predicted CATH superfamily using SSAP. 90.86% of the predictions could be confirmed by structural validation. This is comparable as the CATHe model has an accuracy of 85.6% +− 0.4%, which was obtained for the test set of the TOP 1773 SUPERFAMILIES dataset. We consider this to be a compelling result as Pfam domains do not always correspond to a single domain, they can comprise partial or multiple domains which would make it difficult to meet the strict SSAP score and overlap thresholds that we apply (see supplementary S2.4. for thresholds used).

Furthermore, despite expanding CATH superfamilies with AlphaFold structures brought in via HMM matches, for many superfamilies, less than 20% of the relatives have structural characterization. These characterized relatives may be extremely remote homologues of the proteins in the Pfam-human dataset and therefore difficult to meet the thresholds. In the largest 50 superfamilies in CATH, the structural overlap between very distant homologues can fall below 50%. Once DeepMind releases AlphaFold structures for UniRef90 we will be able to pull more structural data into CATH and re-evaluate whether some of these putative unrelated domains are in fact very remote homologues of CATH superfamilies. Our manual evaluation of the Pfam human matches that did not cross the class-wise SSAP and overlap thresholds showed that the majority (~90%) had the correct protein domain architecture (A-level in CATH), suggesting that CATHe is able to capture important elements of the structural environment.

In summary, our method was tested on a dataset of very remote homologues (less than 20% sequence identity). To our knowledge, it is the only method that has been evaluated on such a strict dataset and subsequently validated by structural comparison using AlphaFold 3D models for predicted classifications. We demonstrated that 4.62 million Pfam domains previously unmapped to CATH could be brought in using a reliable threshold on accuracy and error rate (0.5% error rate). This will expand CATH by 3.06%. However, Pfam does not classify all of UniProt and we will apply CATHe in the future to detect additional domains in UniProt, not classified in Pfam, that can be assigned to CATH superfamilies. We will use good-quality AlphaFold structures to confirm these assignments.

CATHe represents a highly accurate method to classify new protein domains into CATH superfamilies. This gives rise to new avenues for bringing in highly remote homologues from a number of sequence databases. The CATH domain sequence embeddings generated by ProtT5 and the source code for the CATHe model can be found at https://github.com/vam-sin/CATHe.

## 5. Materials and Methods

### 5.1. TOP 1773 SUPERFAMILIES Dataset

The aim of this study is to develop a deep learning tool to detect remote homologues for CATH superfamilies. In order to achieve this, it is necessary to generate a dataset from which the classifiers can learn. We made sure that the testing and validation sets of the dataset consisted only of sequences from the Protein Data Bank (PDB)(2). Whereas the training set contained sequences from CATH-Gene3D (v4.3). CATH-Gene3D contains all the Uniprot sequences which can be assigned to CATH superfamilies via scanning against CATH-HMMs (3) (which uses an e-value of 1e-3). The following steps were conducted to generate the dataset for this task:

a. Cluster the sequence domains present in CATH from the PDB using MMseqs2 (9) with a 20% sequence identity filter. To gain an in-depth understanding of the data processing using MMseqs2, see supplementary section S2.1.
b. Using the MMseqs2 output from step (a), choose those superfamilies that have at least 2 PDB sequences so that they can be split into testing and validation sets. The remaining superfamilies are used to create a “mixed-bag” class summarizing all “Other” superfamilies classes.
c. Build the training set using all the sequence domains from CATH-Gene3D for all the superfamilies decided in step (b) (i.e. superfamilies that have at least 2 PDB S20 sequences, and “Other” superfamilies).
d. Use MMseqs2 to reduce the sequence identity between training and the other two (testing and validation) sets to less than 20%.
e. As a further check, use BLAST to remove homologous sequences, at 20% sequence identity, between the three sets.
f. Randomly under-sample the “Other” superfamilies class to reduce the class imbalance in the dataset.
g. Generate embeddings for the protein sequences using the protein language models (pLMs) ProtBERT and ProtT5 (25). To understand more about the pLMs and the embedding generation process, please refer to the supplementary section S2.1.

After step (a), we identified 1773 CATH superfamilies, having at least two non-redundant PDB sequences. We set the threshold at two sequences because that is the minimum number of sequences that we need in order to split the resultant set into testing and validation sets. The rest of the superfamilies in CATH (3456 superfamilies) were used to make the “Other” superfamilies class. The 1773 superfamilies had a total of 82,720,883 domain sequences in the CATH database (which is 87.6% of the CATH v4.3 database) before being processed. The number of sequences in each of the three sets after processing is given in Table 1. This dataset is referred to as the “TOP 1773 SUPERFAMILIES” dataset.

**Table 1.** Description of the training, validation, and testing sets for the two datasets generated by processing the data from the CATH database.

| Dataset | No. of Training Sequences | No. of Validation Sequences | No. of Testing Sequences |
| --- | --- | --- | --- |
| TOP 1773 SUPERFAMILIES | 1,039,135 | 6,863 | 6,862 |
| TOP 50 SUPERFAMILIES | 528,863 | 1,948 | 1,946 |

### 5.2. TOP 50 SUPERFAMILIES Dataset

To investigate more closely the performance on large (i.e., with high numbers of predicted domains) and highly diverse superfamilies, a subset of the TOP 1773 SUPERFAMILIES dataset which summarizes the 50 largest CATH superfamilies was created. This set holds a total of 39,625,167 domain sequences in the CATH database (37.32% of the CATH domain sequences) and will be dubbed TOP 50 SUPERFAMILIES throughout the text.

The number of sequences in training, testing, and validation sets after processing are given in Table 1.

### 5.3. Human Pfam Domains

There are a total of 36,318 Pfam domains from *homo sapiens* which are not in CATH. In order to study the performance of CATHe and structurally validate those predictions, we decided to choose only those domains which did not have any structurally problematic regions. By problematic, we mean domains with regions that will make it harder to detect structural relationships with CATH structural domains. The structures were obtained from the EBI AlphaFold 2 database (https://alphafold.ebi.ac.uk/). An AlphaFold domain was considered problematic if it met any of the following conditions:

a. Model quality (pLDDT) is less than 90%.
b. Less than three secondary structures in the 3D structure of the domain. This was calculated using DDSP (43).
c. The longest problematic region (the longest stretch of residues that had less than 70% pLDDT) covered more than one-third of the domain length.
d. Disorder greater than 70%. This was also measured using DSSP.

### 5.4. Models

Traditional bioinformatics techniques, as well as advanced machine learning methods, were used to develop classifiers to detect remote homologues. The various techniques that were used are outlined below:

#### BLAST Homology Based Prediction

To develop the BLAST (44) Homology Based Predictor tool, the NCBI BLAST+ toolkit (version 2.11.0) was used. Firstly, a target database was built from the training set using the makeblastdb command. The testing set was used as the query and searched against the target database using the blastp command. For the blastp search, the e-value cutoff was set at 1e+6. This was done in order to obtain hits, even insignificant hits, for all the sequences in the testing set.

Once all the possible hits were obtained for all the sequences in the testing set, they were analyzed to find the most significant hit. For each sequence in the testing set, this was done by first finding the hits with the lowest e-value. Among the hits with the lowest e-value, the hit with the greatest percent sequence identity was chosen. This hit was deemed to be the prediction made by the BLAST model.

#### Artificial Neural Network (ANN)

*A* simple ANN was developed for the task of remote homologue detection. The ANN had one hidden layer consisting of 128 nodes. The hidden layer was followed by the output layer. To reduce overfitting, a dropout (45) layer with a rate of 0.3 and a batch normalization (46) layer were added to the model. The hidden layer used a Rectified Linear Unit (ReLU) (47) activation function, whereas the output layer used a Softmax (48) activation function. The Adam (49) optimizer was used with an initial learning rate of 1e-3. The model was set to train for a maximum of 200 epochs, but early stopping was implemented to prevent the model from overfitting. Early stopping was measured on the validation accuracy with the patience set to 20 epochs. A batch size of 256 was used for the training process. Three ANNs were trained independently on three different features, i.e., ProtBERT embeddings, ProtT5 embeddings, and Protein Sequence Lengths, to make the “ANN + ProtBERT”, “CATHe” (The ANN model trained on the ProtT5 embeddings), and “ANN + Sequence Length” models, respectively.

To arrive at the architecture that we use for the ANN, we conducted an optimization study where we measured the performance of different ANN architectures trained on the ProtT5 embeddings on the testing set, with a lower or higher number of parameters on the two datasets, TOP 1773 SUPERFAMILIES, and TOP50 SUPERFAMILIES. The optimal architecture decided by this study was used to make the models. The details of this study can be found in section S2.2 of the supplementary.

The significance of the “ANN + Sequence Length” model comes from the assumption that the protein sequence embeddings generated by the pLMs, though they were of fixed size, were able to distinguish between small and long sequences. This model was developed in order to make sure that the classification of the protein sequences into CATH superfamilies was not simply based on their respective sequence lengths.

#### Logistic Regression (LR)

Logistic Regression (LR) is a traditional machine learning method that is often used as a baseline to put the performance of more complex ANN models into perspective. In this study, we developed an LR model using the SciKit-Learn (50) python library. An “lgbfs” solver was used with the maximum iterations set to 5,000. The other parameters were set to the default values. This LR model was trained on the ProtBERT and ProtT5 embeddings to make the LR + ProtBERT and LR + ProtT5 models, respectively.

#### Random

The random predictor model assigns an output class for each of the sequences at random taking into account the class imbalance. This serves as an important sanity check for measuring and comparing the performance of the proposed model.

In Fig. 4, we outline the data preprocessing steps and the various models that we have used in this study in order to develop a tool for the detection of remote homologues in the CATH database. Table S7 in the supplementary has a summary of the seven different models developed in this study.

**Fig 4.**
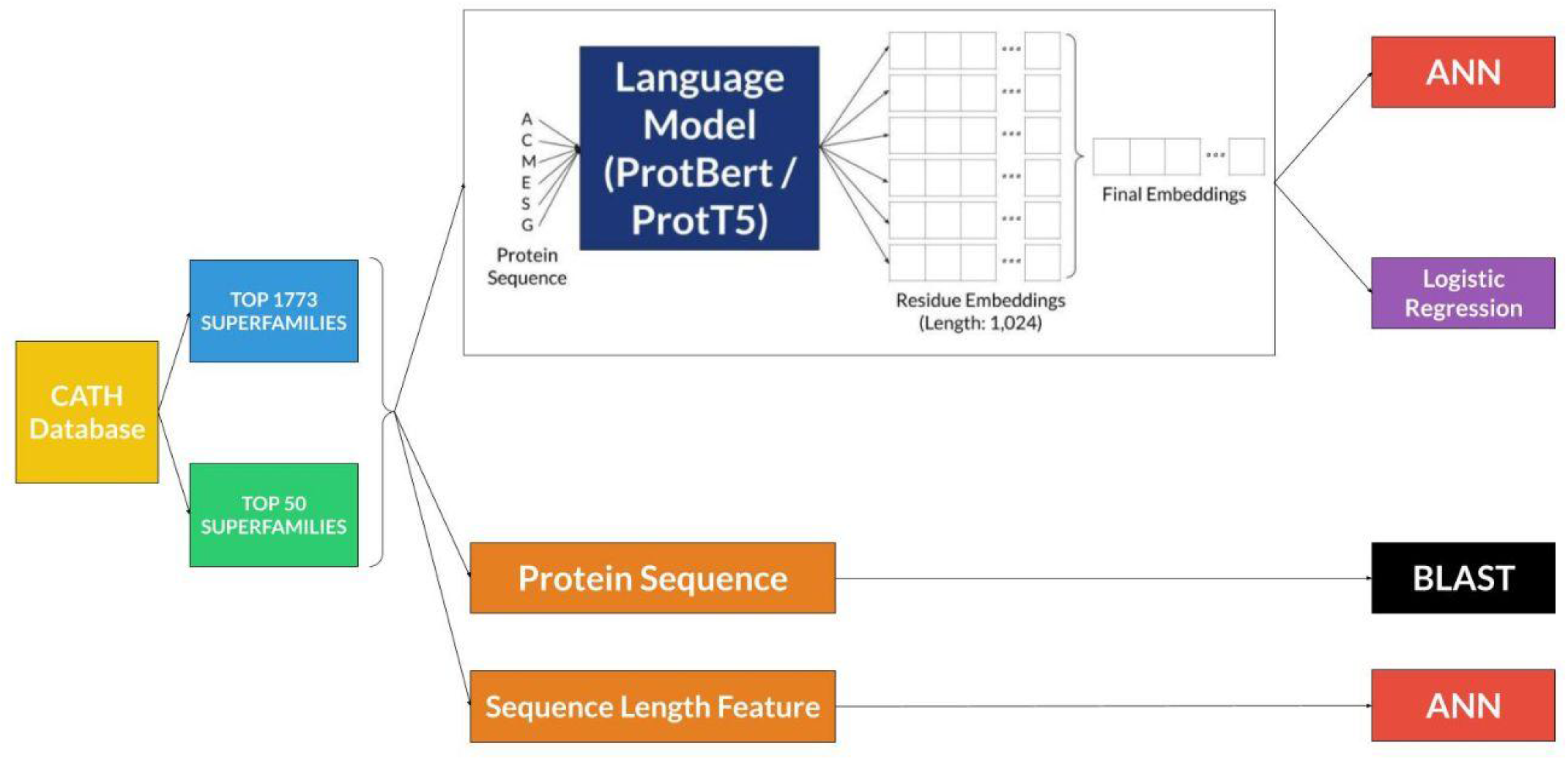
Illustration of the pipeline followed in the study which includes the generation of the two datasets, TOP 1773 SUPERFAMILIES and the TOP 50 SUPERFAMILIES, the features extracted from these datasets, and the various models that were developed.

### 5.5. Metrics

Four different metrics, accuracy, F1-Score, MCC, and balanced accuracy, were used to measure the performance of the models in this study. Please refer to section S2.3. in the supplementary material for more information on the metrics and how they are calculated.

## 6. Data Availability

The source code along with the datasets used for training are available at https://github.com/vam-sin/CATHe

## 7. Acknowledgments

We would sincerely like to thank David Gregory and Tristan Clarke from the Research Computing team at University College London.

