## Supplementary Document for "CATHe: Detection of remote homologues for CATH superfamilies using embeddings from protein language models"

**Outline:**

This supplementary document provides an in-depth analysis of the CATHe model and explains in detail the various parts of the development pipeline, such as dataset preprocessing, and model optimization. Additionally, the different experiments conducted to decide the structural validation thresholds and CATHe prediction probability thresholds are discussed.

### S1. Results

##### S1.1. TOP 1773 SUPERFAMILIES

| Model | Accuracy | F1-Score | MCC | Bal Acc |
|---|---|---|---|---|
| ANN + ProtBERT | 74.42 +- 0.53 | 54.47 +- 0.71 | 0.7434 +- 0.0053 | 58.92 +- 0.67 |
| LR + ProtBERT | 61.31 +- 0.59 | 34.39 +- 0.62 | 0.6115 +- 0.0059 | 37.91 +- 0.62 |
| **CATHe** | **85.60 +- 0.41** | **72.35 +- 0.67** | **0.8554 +- 0.0041** | **76.11 +- 0.62** |
| LR + ProtT5 | 83.11 +- 0.44 | 66.96 +- 0.72 | 0.8309 +- 0.0044 | 70.3 +- 0.67 |
| ANN + Length | 3.33 +- 0.21 | 0.04 +- 0.004 | 0.0273 +- 0.0023 | 0.23 +- 0.01 |
| BLAST | 31.24 +- 0.56 | 21.53 +- 0.60 | 0.31 +- 0.0056 | 25.31 +- 0.66 |
| Random | 0.44 +- 0.08 | 0.04 +- 0.02 | 0.00003 +- 0.00080 | 0.07 +- 0.03 |

**Table S1 (A). Performance of seven different models, ANN+ProtBERT, LR+ProtBERT, CATHe, LR+ProtT5, ANN+Sequence Length, BLAST, and Random on the TOP 1773 SUPERFAMILIES dataset, measured using four different metrics, Accuracy, F1-Score, MCC, and Balanced Accuracy along with 95% confidence intervals**

| Model | Accuracy | F1-Score | MCC | Bal Acc |
|---|---|---|---|---|
| ANN + ProtBERT | 94.39 +- 0.51 | 91.78 +- 1.07 | 0.9417 +- 0.0053 | 92.72 +- 0.8 |
| LR + ProtBERT | 88.86 +- 0.71 | 86.25 +- 1.23 | 0.8844 +- 0.0073 | 86.39 +- 1.13 |
| **CATHe** | **98.15 +- 0.30** | **95.49 +- 0.93** | **0.9807 +- 0.0031** | **95.98 +- 0.67** |
| LR + ProtT5 | 97.73 +- 0.34 | 95.58 +- 0.67 | 0.9764 +- 0.0035 | 95.44 +- 0.71 |
| ANN + Length | 11.00 +- 0.69 | 3.13 +- 0.19 | 0.0974 +- 0.0079 | 5.29 +- 0.31 |
| BLAST | 53.90 +- 1.12 | 49.77 +- 1.42 | 0.5266 +- 0.0114 | 57.84 +- 1.54 |

| | | | | |
|---|---|---|---|---|
| Random | 2.68 +- 0.36 | 1.60 +- 0.28 | 0.00001 +- 0.00030 | 2.04 +- 0.57 |

**Table S1 (B). Performance of seven different models, ANN+ProtBERT, LR+ProtBERT, CATHe, LR+ProtT5, ANN+Sequence Length, BLAST, and Random on the TOP50 SUPERFAMILIES dataset, measured using four different metrics, Accuracy, F1-Score, MCC, and Balanced Accuracy along with 95% confidence intervals**

The performance of the seven prediction models, BLAST, ANN + ProtBERT, CATHe, and ANN + Sequence Length, LR + ProtBERT, LR + ProtT5, and Random, was measured on the testing set of the TOP 1773 SUPERFAMILIES dataset using four metrics, Accuracy, F1-Score, MCC, and Balanced Accuracy. The results have been illustrated in Fig. 1 (A) and described in Table S1 (A).

The CATHe model had the greatest performance in terms of all the four metrics, with an accuracy of 85.60% +- 0.41%, an F1-Score of 72.35% +- 0.67%, an MCC of 0.8554 +- 0.0041, and a Balanced Accuracy of 76.11% +- 0.62%. In terms of the pLM embeddings, we noticed that the ProtT5 embeddings led to greater performance compared to ProtBERT embeddings (Fig 1 (A)). CATHe had a much greater performance compared to the BLAST model applying homology-based inference. Additionally, from the poor performance of the ANN + Sequence Length model, we can infer that the sequence length is not a distinguishing feature that can be used to classify the protein sequences into CATH superfamilies.

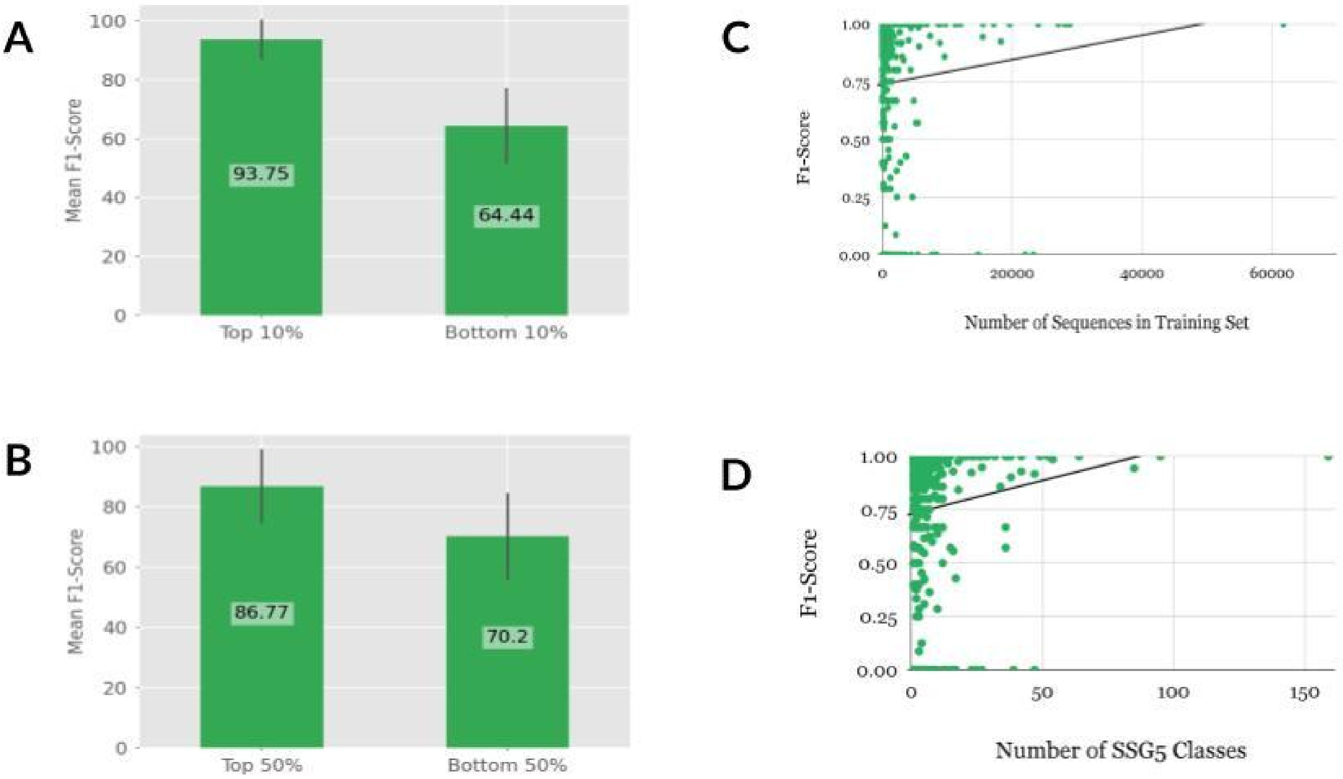

**Fig S1. Analysis of the CATHe model on the TOP 1773 SUPERFAMILIES dataset. (A) Mean F1-Score of the CATHe model on the testing sequences of the superfamilies consisting of the top 10% and bottom 10% of the training dataset of the TOP 1773 SUPERFAMILIES dataset. (B) Mean F1-Score of the CATHe model on the testing sequences of the superfamilies consisting of the top 50% and bottom 50% of the training dataset of the TOP 1773 SUPERFAMILIES dataset. (C) Superfamily population in terms of the number of sequences present in the training set for that superfamily measured against the F1-Score obtained by CATHe on the testing sequences of that superfamily. (D) F1-Score obtained by the CATHe on the testing sequences of a superfamily measured against the number of Structurally Similar Groups (SSG5) classes present in the superfamily.**

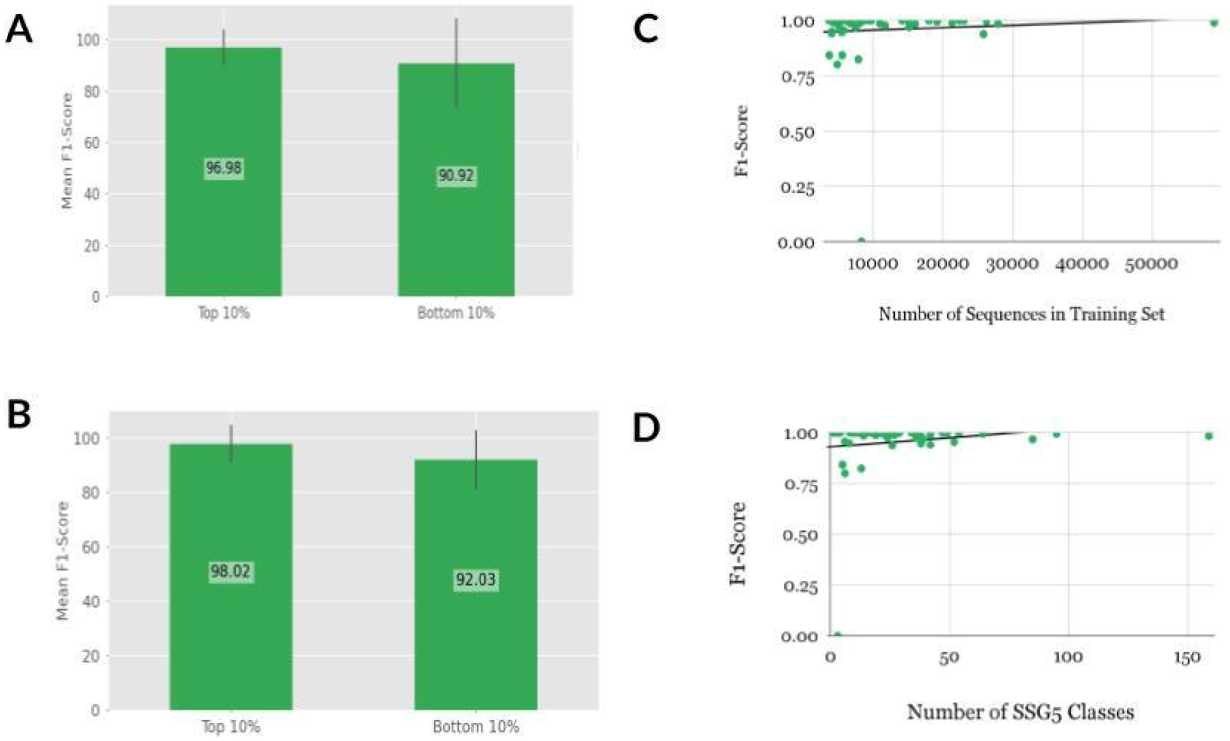


**Fig S2. Analysis of the CATHe model trained on the TOP50 SUPERFAMILIES dataset. (A) Mean F1-Score of the CATHe model on the testing sequences of the superfamilies consisting of the top 10% and bottom 10% of the training dataset of the TOP50 SUPERFAMILIES dataset. (B) Mean F1-Score of the CATHe model on the testing sequences of the superfamilies consisting of the top 50% and bottom 50% of the training dataset of the TOP50 SUPERFAMILIES dataset. (C) Superfamily population in terms of the number of sequences present in the training set for that superfamily measured against the F1-Score obtained by the CATHe on the testing sequences of that superfamily. (D) F1-Score obtained by the CATHe on the testing sequences of a superfamily measured against the number of Structurally Similar Groups (SSG5) classes present in the superfamily.**

To understand how the imbalance in the training set of the TOP 1773 SUPERFAMILIES, and TOP 50 SUPERFAMILIES datasets affects the performance of CATHe on individual superfamilies, we conducted four comparisons.

1. We compared the mean F1-Score on the superfamilies consisting of the top 10%, and the bottom 10% of the training set. To make the top 10%, and bottom 10%, the superfamilies were first organized in the descending order of their population in the training set. For the top 10%, we chose the first X number of superfamilies in the descending order that can together make up 10% of the training set in terms of the number of sequences. Similarly for the bottom 10%, we chose the last Y number of superfamilies that can together make up 10% of the training set in terms of the number of sequences.

   For the TOP 1773 SUPERFAMILIES, the mean F1-Score on the top 10% superfamilies (which is the 3 largest superfamilies) was almost 29 percentage points more than the mean F1-Score on the bottom 10% superfamilies (which is the 1267 smallest superfamilies). This significant drop in performance indicates how greatly the number of sequences for a superfamily in the training set affects the performance of CATHe (Fig S1 (A)).

   For the TOP 50 SUPERFAMILIES, we notice a similar trend. The mean F1-Score on the superfamilies consisting of the top 10% (which is the largest superfamily), is greater than that on the bottom 10% (which is the 13 smallest superfamilies) by almost 3 percentage points bottom 10% (Fig S2 (A)).

2. Comparison of the mean F1-Score on the superfamilies consisting of the top 50%, and the bottom 50% of the training set. The top 50%, and the bottom 50% are made in the same fashion as the top 10%, and bottom 10%.

   For the TOP 1773 SUPERFAMILIES dataset, there is a decrease in performance, a difference of almost 16 percentage points, from the mean F1-Score on the top 50% (which is the 47 largest superfamilies) and the bottom 50% (which is the 1727 smallest superfamilies) superfamilies (Fig S1 (B)). Although, this difference in performance is not significant.

   The result for the TOP 50 SUPERFAMILIES is similar, the mean F1-Score on the superfamilies consisting of the top 50% (which is the 11 largest superfamilies), is greater than that on the bottom 50% (which is the 39 smallest superfamilies) by almost 4 percentage points. (Fig S2 (B))

3. A broader analysis was conducted by making a scatter plot comparing the F1-Score of the CATHe model on a superfamily with the population of the superfamily (in terms of the number of sequences) in the training set. Fig S1 (C) (TOP 1773 SUPERFAMILIES) and Fig S2 (C) (TOP 50 SUPERFAMILIES) compare on an individual superfamily level, how the

population of a superfamily in terms of the number of sequences in the training set, affects the CATHe performance. For the TOP 1773 SUPERFAMILIES, and the TOP 50 SUPERFAMILIES, the Pearson Correlation Coefficient (r) for these two features is 0.0983, and 0.077 respectively, which means there is no (or negligible) direct linear relationship between the two features in both these cases.

4. In Fig. S1 (D) and Fig S2 (D), we plot the performance on individual superfamilies, in terms of F1-Score, against the number of Structurally Similar Groups at 5 Å threshold (SSG5 classes) present in that superfamily for the TOP 1773 SUPERFAMILIES and the TOP 50 SUPERFAMILIES dataset respectively.

   Each superfamily is clustered into smaller groups defined by their structural similarity. SSG5 classes have a threshold of 5 Å threshold for their Root Mean Square Deviation (RMSD). The number of SSG5 classes in a superfamily can be used as a measure of structural diversity, a greater number of SSG5 classes in a superfamily would mean greater structural diversity in that superfamily.

   This plot was done in an effort to understand how structural diversity affects the performance of the model on that superfamily. The Pearson Correlation coefficient (r) for the performance (F1-Score) and the structural diversity (Number of SSG5 classes) came out to be 0.054, and 0.171 for the TOP 1773 SUPERFAMILIES, and the TOP 50 SUPERFAMILIES respectively, which means there is no (or negligible) direct linear relationship between the two features in both these cases.

***Threshold Analysis on TOP 1773 SUPERFAMILIES dataset***

In order to use the CATHe model to make predictions on new domains, and to obtain a measure of the reliability of these predictions, we measured the performance (in terms of accuracy, precision, recall, and F1-Score) of the model on the validation set (consisting of 6,863 domains) of the TOP 1773 SUPERFAMILIES dataset at various prediction probabilities. The results from this analysis have been described in Table S2 and Figure S2. Using a prediction probability threshold of 0.9, we obtain an optimal performance (99.5% accuracy / 0.5% error rate) without compromising on the number of domain predictions that cross the threshold (3415/6863; 49.75% coverage of the validation set) (Table S2).

| **Threshold** | **Accuracy** | **Precision** | **Recall** | **F1-Score** | **Number of predictions that cross the threshold (coverage)** |
|---|---|---|---|---|---|
| 0.99 | 99.70 | 98.14 | 98.14 | 98.14 | 344 |

| | | | | | |
|---|---|---|---|---|---|
| 0.98 | 99.66 | 98.27 | 98.59 | 98.38 | 907 |
| 0.97 | 99.78 | 98.70 | 98.97 | 98.79 | 1405 |
| 0.96 | 99.67 | 98.62 | 98.91 | 98.74 | 1848 |
| 0.95 | 99.64 | 98.82 | 99.04 | 98.91 | 2242 |
| **0.9** | **99.50** | **98.14** | **98.38** | **98.18** | **3415** |
| 0.85 | 99.44 | 98.30 | 98.52 | 98.34 | 4140 |
| 0.8 | 99.23 | 97.76 | 98.10 | 97.83 | 4553 |
| 0.75 | 99.08 | 97.38 | 97.73 | 97.44 | 4827 |
| 0.7 | 98.90 | 97.06 | 97.48 | 97.12 | 5045 |
| 0.65 | 98.62 | 96.51 | 96.99 | 96.57 | 5240 |
| 0.6 | 98.38 | 96.16 | 96.63 | 96.19 | 5386 |
| 0.55 | 98.02 | 95.30 | 95.82 | 95.29 | 5510 |
| 0.5 | 97.65 | 94.66 | 95.16 | 94.60 | 5627 |
| 0.45 | 97.15 | 93.20 | 94.03 | 93.27 | 5720 |
| 0.4 | 96.70 | 92.18 | 93.26 | 92.31 | 5820 |
| 0.35 | 96.08 | 91.05 | 92.22 | 91.15 | 5924 |
| 0.3 | 95.29 | 88.97 | 90.35 | 89.13 | 6033 |
| 0.25 | 94.11 | 86.26 | 87.99 | 86.46 | 6149 |
| 0.2 | 92.94 | 83.82 | 85.77 | 84.04 | 6266 |
| 0.15 | 91.75 | 80.99 | 83.25 | 81.24 | 6395 |
| 0.1 | 89.83 | 77.54 | 79.95 | 77.71 | 6570 |
| 0.05 | 87.07 | 72.99 | 75.45 | 73.03 | 6800 |
| 0 | 86.33 | 71.89 | 74.19 | 71.82 | 6863 |

**Table S2. Performance of CATHe, measured in terms of accuracy, precision, recall, and f1-score, on the validation set of the TOP 1773 SUPERFAMILIES dataset at different prediction probability thresholds.**

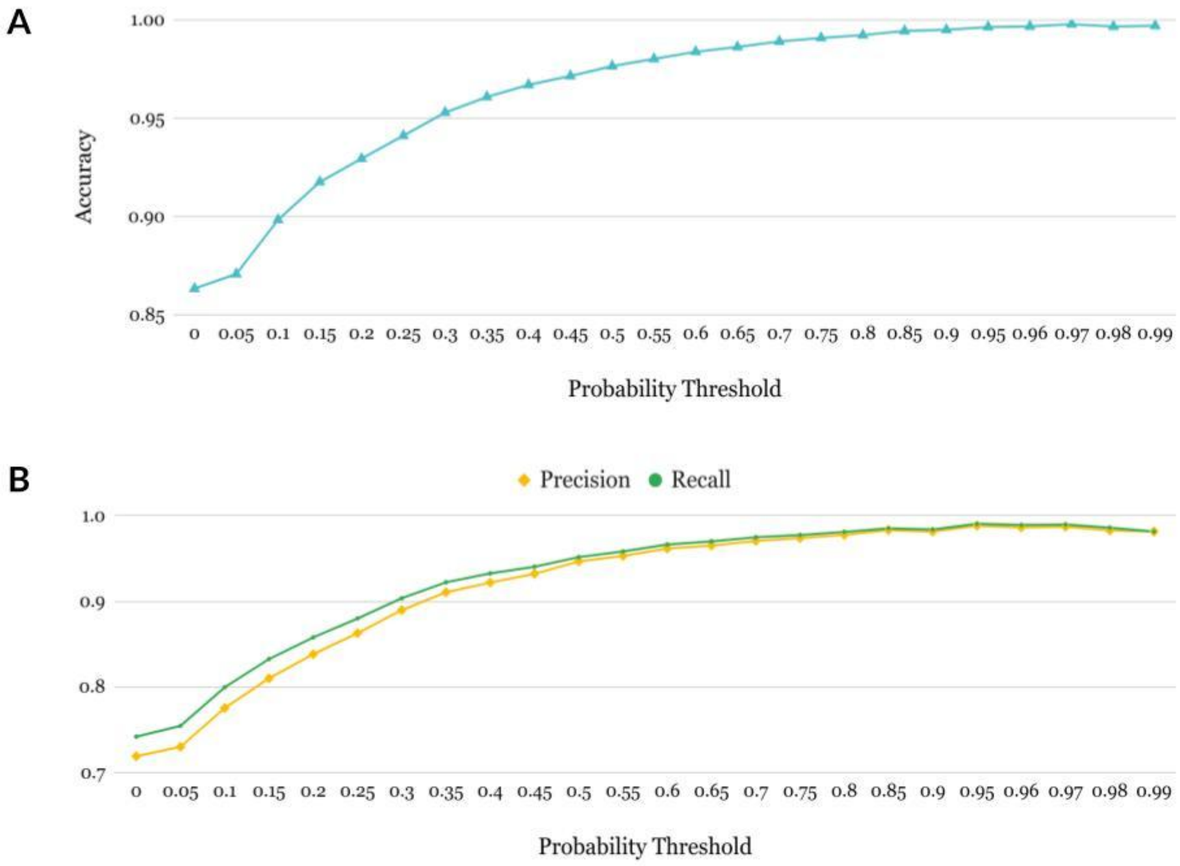


**Figure S3. Performance of CATHe on the validation set of the TOP 1773 SUPERFAMILIES dataset at different prediction probability thresholds. Panel (A) shows the accuracy at various probability thresholds; panel (B) the precision and recall.**

***Misclassification analyses on the TOP 50 SUPERFAMILIES dataset***

Additionally, we analyzed all the misclassifications made by the CATHe model. There were a total of 36 misclassifications on the testing set of the TOP50 SUPERFAMILIES dataset (Table S1). Out of the 36 errors in the Hierarchical Superfamily, 28 are errors in Topology, 22 are errors in Architecture, and 13 are errors in Class.

| CATH Domain | Ground Truth Superfamily Annotation | Predicted Superfamily | Prediction Probability |
|---|---|---|---|
| 2lvsA01/1-56 | 1.10.10.60 | 1.10.10.10 | 99.99% |
| 2ipqX01/396-417_439-471 | 1.10.10.10 | 1.10.150.240 | 47.52% |

| | | | |
|---|---|---|---|
| 1ef4A00/1-55 | 1.10.10.60 | 1.10.10.10 | 67.68% |
| 2nn6H01/25-79 | 2.40.50.100 | 3.50.50.60 | 16.11% |
| 2w48B01/3-54 | 1.10.10.60 | 1.10.10.10 | 62.22% |
| 3tw6A06/950-1003 | 1.10.10.60 | 3.20.20.70 | 79.08% |
| 2f93B00/29-79 | 1.10.287.470 | 1.20.1560.10 | 44.43% |
| 2nn6I01/6-56 | 2.40.50.100 | 2.40.50.140 | 36.25% |
| 1rsoA01/7-57 | 1.10.287.470 | 1.10.10.10 | 21.07% |
| 1pryA01/2-51 | 3.30.200.20 | 2.40.50.100 | 58.48% |
| 3bg3A04/982-1028 | 1.10.10.60 | 3.20.20.70 | 69.82% |
| 1h6lA00/29-381 | 2.120.10.30 | 2.130.10.10 | 80.34% |
| 2r1fB01/81-109 | 3.30.160.60 | 2.40.50.140 | 66.89% |
| 1k32A02/326-675 | 2.130.10.10 | 2.120.10.30 | 73.73% |
| 3brcB01/2-35 | 1.10.287.470 | 1.10.150.240 | 38.16% |
| 1fcqA00/10-330 | 3.20.20.70 | 3.20.20.80 | 94.84% |
| 4fnsA02/310-632 | 3.20.20.70 | 3.20.20.80 | 94.00% |
| 1ia9A01/1576-1722 | 3.30.200.20 | 1.25.40.10 | 50.14% |
| 4pz8A01/2-5_244-373 | 2.40.50.140 | 3.40.50.300 | 31.24% |
| 1ev7B02/177-309 | 1.10.10.10 | 2.40.50.140 | 81.81% |
| 2yhxA02/59-188 | 3.30.420.40 | 3.40.190.10 | 49.93% |
| 4nrtA03/216-251_301-387 | 3.30.70.270 | 3.20.20.70 | 44.10% |
| 2eg9A02/121-147_204-283 | 3.40.50.720 | 2.60.120.260 | 19.67% |
| 5oomW00/40-148 | 2.40.50.100 | 2.40.50.140 | 88.21% |
| 3c5mA00/2-387 | 2.130.10.10 | 2.120.10.30 | 62.18% |
| 2ea9A01/17-110 | 3.30.450.20 | 3.40.50.300 | 14.35% |
| 1ckmA02/236-318 | 2.40.50.140 | 3.40.190.10 | 21.25% |
| 2q7nA03/204-283 | 2.60.40.10 | 3.40.50.300 | 93.52% |
| 5d6eA02/372-451 | 1.10.10.10 | 1.10.150.240 | 24.32% |
| 5gjkA01/461-535 | 1.10.10.10 | 3.40.50.300 | 84.73% |
| 1d3yA01/72-142 | 1.10.10.10 | 1.10.150.240 | 68.93% |
| 1cidA02/107-177 | 2.60.40.10 | 2.40.50.100 | 94.38% |

| | | | |
|---|---|---|---|
| 2r7dA01/3-71 | 1.10.10.10 | 1.10.150.240 | 93.88% |
| 2imrA01/34-45_51-90_400-416 | 2.30.40.10 | 2.60.120.10 | 26.46% |
| 1v8qA00/20-85 | 2.40.50.100 | 2.40.50.140 | 70.88% |
| 2nyvA02/18-81 | 1.10.150.240 | 3.30.420.40 | 62.07% |

**Table S3. Misclassifications made the CATHe model on the testing set of the TOP50 SUPERFAMILIES dataset.**

*T-SNE Analysis on the TOP50 Superfamilies Dataset*

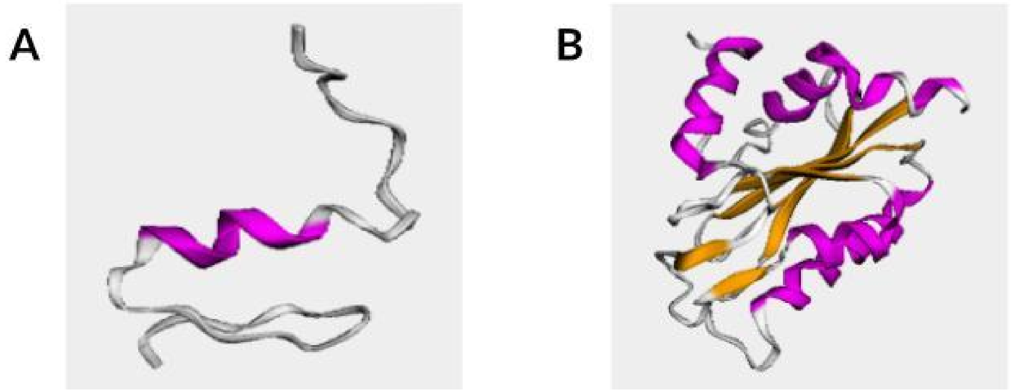


**Fig S4. Domains from the 3.30 CATH architecture. (A) Domain 2enhA01 from the 3.30.160.60 superfamily. (B) Domain 3bp8A02 from the 3.30.420.40 superfamily.**

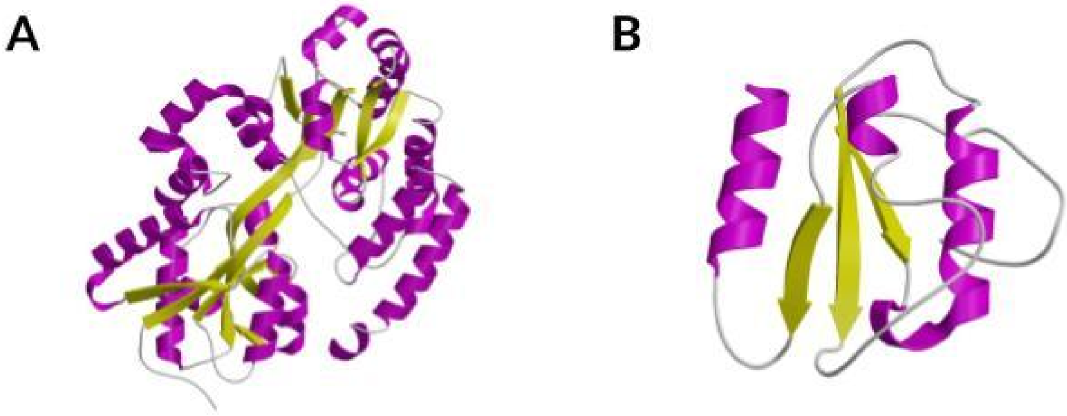


**Fig S5. Domains from the 3.40 CATH architecture. (A) Domain 4mfiA00 from the 3.40.190.10 superfamily. (B) Domain 4c0lA03 from the 3.40.50.300 superfamily.**

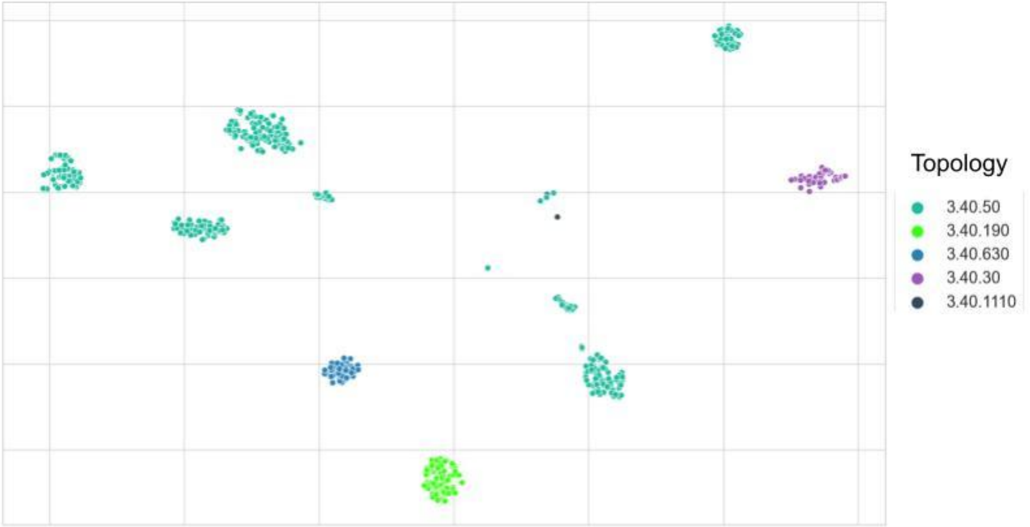


**Fig S6. T-SNE projections of high-dimensional embedding spaces for the TOP50 SUPERFAMILIES test set colored by CATH architectures. The embeddings were extracted from the last layer of the trained CATHe ANN to highlight the additional benefit of supervised training for distinguishing CATH architectures. In this figure, we show the embeddings for domains belonging to the various fold groups in the 3.40 architecture.**

***Human Pfam Domains Analysis***

In the Human-Pfam set there were 197 domains, out of which 142 crossed the individual class-wise thresholds for SSAP score and Structural Overlap percentage. We conducted manual curation on the 55 domains that did not cross the thresholds. We analyzed which level of the CATH hierarchy (Class, Architecture, Topology, and Homologous Superfamily) the CATHe match belonged to and tabulated the results in Table S4. Through the manual curation we noticed that out of the 55 domains, 37 matched at the superfamily level, whereas 4 match at the topology level, 9 matched at the architecture level, and 5 matched at the class level. This would bring the total domains matched at the superfamily level to 179 out of 197 (90.86%).

| **CATHe predicted superfamily** | **Protein 1: Human Pfam Domain** | **Protein 2: CATH domain from AlphaFold expanded superfamilies** | **SSAP Score** | **Structural Overlap Percentage** | **Lowest match level: C, A, T, H** |
|---|---|---|---|---|---|
| 1.10.10.10 | af_Q9Y2G1_587_647 | af_P9WKH5_4_69 | 70.84 | 51 | H |
| 1.10.1070.11 | af_Q96MK3_305_522 | af_A4I109_891_1081 | 69.43 | 71 | H |

| | | | | | |
|---|---|---|---|---|---|
| 1.20.1250.20 | af_Q9H1C4_115_194 | af_Q54M69_1_130 | 74.67 | 57 | H |
| 1.20.1250.20 | af_Q9HAB3_273_371 | af_A1Z9U2_489_682 | 70.92 | 45 | H |
| 1.20.1250.20 | af_Q9NQ40_297_395 | af_A0A0G2L0H0_282_469 | 70.8 | 47 | H |
| 1.20.144.10 | af_Q8NHU3_220_293 | af_Q54FF5_14_140 | 69.36 | 57 | A |
| 1.20.144.10 | af_Q96LT4_292_365 | af_Q54FF5_14_140 | 69.4 | 47 | H |
| 1.25.10.10 | af_Q8N612_96_426 | af_I1K1L7_3_208 | 71.75 | 53 | H |
| 1.25.10.10 | af_Q9NSG2_176_728 | af_A0A2R8PW20_341_915 | 64.73 | 67 | H |
| 1.25.10.10 | af_Q9ULQ0_49_324 | af_A0A1D6LVL4_460_650 | 75.03 | 59 | H |
| 1.25.10.10 | af_Q9Y3T9_327_622 | af_Q54N00_37_333 | 70.87 | 67 | H |
| 1.10.1070.11 | af_O75063_187_399 | af_Q4CY12_25_225 | 69.99 | 68 | A |
| 1.20.1230.10 | af_O94804_755_895 | af_P13217_853_1091 | 80.33 | 57 | A |
| 1.25.10.10 | af_O95155_591_1212 | af_P35222_123_666 | 66.41 | 54 | A |
| 1.25.10.10 | af_Q14997_330_828 | af_P25339_554_886 | 67.59 | 61 | H |
| 1.25.40.10 | af_Q15573_63_423 | af_F1LRQ6_328_538 | 73.67 | 55 | H |
| 1.25.10.10 | af_Q68E01_271_495 | af_B3CJ34_1_236 | 68.31 | 78 | A |
| 1.20.144.10 | af_Q86VZ5_276_349 | af_Q54FF5_14_140 | 69.68 | 55 | C |

| | | | | | |
|---|---|---|---|---|---|
| 1.10.1070.11 | af_Q8IXL6_353_570 | af_A4I109_891_1081 | 69.56 | 71 | C |
| 2.40.33.20 | af_Q5VT66_202_333 | af_P95151_15_247 | 73.81 | 47 | H |
| 2.60.40.10 | af_Q969Y0_73_284 | af_A0A286Y9T4_97_203 | 82.24 | 50 | H |
| 2.40.33.20 | af_Q969Z3_201_332 | af_P95151_15_247 | 73.63 | 47 | H |
| 2.60.200.20 | af_Q96FA3_8_418 | af_A0A1D8PJZ1_1_154 | 64.83 | 30 | T |
| 2.60.120.10 | af_Q8NE79_40_267 | af_A0A1D6G3C8_362_502 | 78.52 | 59 | H |
| 2.60.200.20 | af_Q9HAT8_10_420 | af_A0A1D8PJZ1_1_154 | 64.77 | 30 | T |
| 2.60.120.10 | af_Q9HBU9_25_251 | af_A0A1D6G3C8_362_502 | 79.45 | 59 | H |
| 2.130.10.10 | af_Q9UKN8_63_254 | af_Q8K1S1_252_395 | 72.37 | 50 | H |
| 3.30.40.10 | af_Q92622_737_937 | af_O22727_1_134 | 67.28 | 49 | C |
| 3.40.960.10 | af_Q969Z0_565_620 | af_Q8IIC7_597_695 | 83.06 | 56 | H |
| 3.40.50.1220 | af_Q9BPY3_157_301 | af_I1KYK3_1_224 | 65.7 | 45 | T |
| 3.90.190.10 | af_Q9C0I1_300_501 | af_A0A0R4IFR5_129_524 | 79.66 | 48 | H |
| 3.40.50.1110 | af_Q9H1Q7_18_171 | af_P0ADA1_24_208 | 71.53 | 48 | T |
| 3.20.20.70 | af_Q9H9T3_312_393 | af_Q58692_136_357 | 63.97 | 33 | H |
| 3.40.50.150 | af_Q9HCE5_186_363 | af_P28638_12_288 | 72.54 | 57 | H |
| 3.40.50.1220 | af_Q9NWS6_142_287 | af_P17109_212_337 | 66.24 | 76 | C |

| | | | | | |
|---|---|---|---|---|---|
| 3.90.190.10 | af_Q9NXD2_328_512 | af_A0A0R4IFR5_129_524 | 79.39 | 45 | H |
| 3.10.110.10 | af_Q9NXR7_8_333 | af_O94721_1_121 | 65.85 | 33 | H |
| 3.40.30.10 | af_Q9P2K2_533_722 | af_Q10057_226_347 | 78.68 | 57 | H |
| 3.40.630.30 | af_Q8NHU2_15_268 | af_P46854_2_162 | 74.12 | 53 | A |
| 3.90.190.10 | af_A4FU01_305_483 | af_A0A0R4IFR5_129_524 | 79.14 | 44 | H |
| 3.90.550.10 | af_O95461_473_540 | af_O05154_2_177 | 63.85 | 35 | H |
| 3.40.30.10 | af_P13284_63_166 | af_I1LLR5_19_195 | 82.44 | 56 | H |
| 3.40.50.2000 | af_P13807_31_663 | af_Q55GH4_22_330 | 78.94 | 43 | H |
| 3.40.50.880 | af_P39656_47_456 | af_P00904_2_193 | 65.27 | 45 | H |
| 3.40.50.2000 | af_P54840_32_667 | af_Q55GH4_22_330 | 78.64 | 43 | H |
| 3.40.960.10 | af_Q14CZ7_593_650 | af_Q84MH1_535_634 | 84.76 | 58 | H |
| 3.40.960.10 | af_Q53R41_779_838 | af_Q84MH1_535_634 | 84.49 | 58 | H |
| 3.30.40.10 | af_Q6ZN54_295_497 | af_O22727_1_134 | 65.37 | 49 | A |
| 3.30.40.10 | af_Q6ZWE6_529_730 | af_O22727_1_134 | 67.28 | 50 | A |
| 3.40.960.10 | af_Q7L8L6_699_758 | af_Q8IIC7_597_695 | 81.81 | 58 | H |
| 3.40.50.150 | af_Q86U44_389_550 | af_P28638_12_288 | 73.07 | 54 | H |
| 3.40.50.150 | af_Q8N0W3_95_496 | af_A0A1D6HWJ4_65_382 | 69.22 | 50 | C |

| | | | | | |
|---|---|---|---|---|---|
| 3.40.50.12780 | af_Q8N2G8_186_506 | af_P31686_1_188 | 63.82 | 46 | A |
| 3.90.550.10 | af_Q8N3Y3_431_498 | af_O05154_2_177 | 64.38 | 35 | H |

**Table S4. Description of the manual curation conducted on the Human Pfam domains which did not cross the SSAP score and Structural Overlap percentage thresholds decided by our threshold analysis. The analysis consisted of manually verifying which level of the CATH hierarchy (Class (C), Architecture (A), Topology (T), and Homologous Superfamily (H)) the CATHe prediction for the domain belonged to.**

### S2. Materials and Methods

#### S2.1. Dataset Processing:

a) **TOP 1773 SUPERFAMILIES dataset:**

***Sequence Redundancy Removal***

The sequence identity filter of 20% we chose in step (a) (refer to Section 5.1.1 in the main manuscript) was obtained from a study (1) on protein sequence and structure relationships at various sequence identity thresholds. According to the study, for sequence identity greater than 30%, structural similarity could be inferred. According to the same study, in the sequence identity range of 20-28%, the majority of residue exchanges that resulted in stable structures resulted in proteins belonging to different functional families. Hence, for the task of detecting remote homologues, we decided on the 20% sequence identity cutoff.

Additionally, in steps (a) & (d) we noticed that sequence clustering tools in general, including MMseqs2, can miss some relationships. This is because these tools result in a high number of false negatives in the twilight region (pairwise sequence identity of 20-35%). To tackle this issue, we double-checked by recursively searching the output sequences against each other using MMseqs2. This way we removed any sequences that might have mistakenly been included in the dataset. As an additional check, in step (e) we use BLAST to remove any leftover homologues. This level of rigorous redundancy removal was conducted to be absolutely sure that all the sequences had less than 20% sequence identity to each other. If this was not the case, then developed models would not be suitable for remote homologue detection.

***Protein Language Model Embeddings***

In this study, we follow a feature-based approach. A feature-based approach is when representations from a language model are generated to be used for training task-specific deep learning architectures. To generate the representations in step (f) (refer to Section 5.1.1 in the main manuscript), we choose to use two protein Language Models (pLMs). The first is ProtBERT, which was developed by training the Bidirectional Encoder Representations from Transformers (BERT) (2) model on the BFD dataset (3). The BERT model relies on a "masked language model" (MLM). The MLM randomly masks some of the input tokens, and the goal of the model is to use only the context to predict the original vocabulary id.

The second pLM we used was ProtT5, which was developed by training the T5 (4) model on the BFD dataset and refined using UniRef50. The T5 model consists of an encoder that projects the source language into an embedding space, and a decoder that takes the embeddings and converts them to the target language. The reason we chose ProtBERT and ProtT5 for our study is that these two pLMs showed the greatest performance in terms of classifying protein sequences into SCOPe structural classes (5).

The following steps were followed to generate the protein sequence embeddings from the pLMs:

- Provide the protein FASTA sequence as the input to the above-described pLMs. These models generate an array of 1,024 values for each of the residues in the input sequence. This output is referred to as the residue-level embeddings.
- A mean-pool operation is conducted on the residue-level embeddings to generate the final protein sequence embedding.

Mean-pooling of the residue-level embeddings, as compared to other operations such as min-pooling, max-pooling, and concatenation, had the greatest performance on a variety of protein sequence level classification tasks (5), hence the mean-pooling operation was preferred. The pLMs were used as static feature encoders (no gradient backpropagation to the pLM).

**b) TOP50 SUPERFAMILIES dataset:**

Fig S7 illustrates the population of each of the superfamilies in the TOP50 SUPERFAMILIES dataset. This population is represented in terms of the percentage of the training set they occupy. The largest, which is 1.25.40.10, occupies 11.13% of the training set, whereas most others occupy less than 2%.

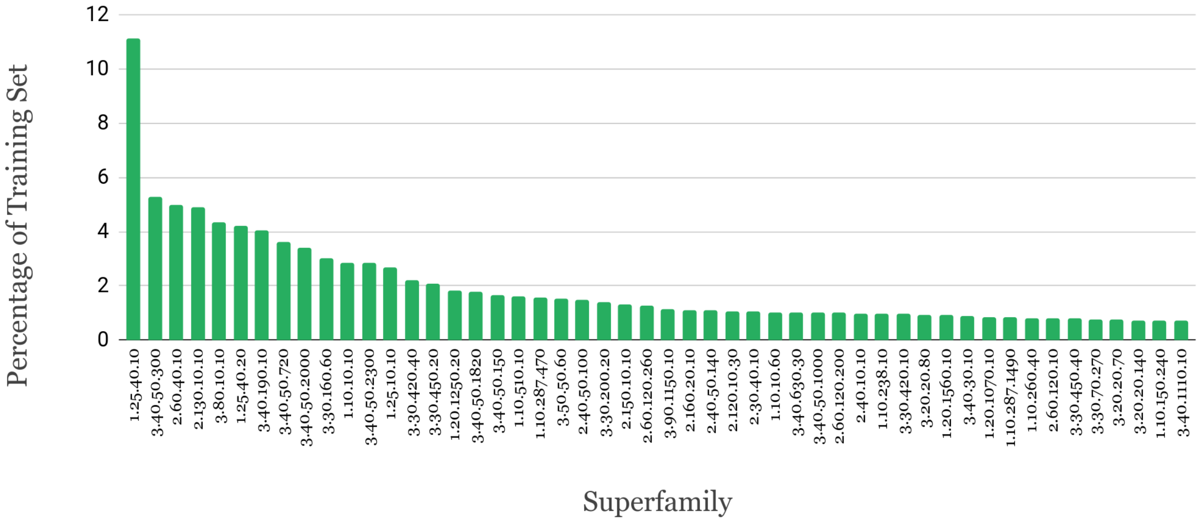


**Fig S7. Distribution of superfamilies in terms of percentage of sequences in the training set for the TOP50 SUPERFAMILIES dataset.**

| Architecture | CATH Superfamily | No. of Sequences (20% PIDE) | No. of SSG5 classes | Sequence Length in Training Set (Mean ± STD) |
|---|---|---|---|---|
| **1.10: Orthogonal Bundle** | 1.10.287.1490 | 4555 | 1 | 161 +- 25 |
| | 1.10.260.40: lambda repressor-like DNA-binding domains | 4319 | 11 | 69 +- 20 |
| | 1.10.150.240: Putative phosphatase; domain 2 | 3704 | 4 | 62 +- 18 |
| | 1.10.510.10: Transferase(Phosphotransferase) domain 1 | 8581 | 46 | 118 +- 63 |
| | 1.10.10.10: Winged helix-like DNA-binding domain superfamily/Winged helix DNA-binding domain | 15123 | 84 | 69 +- 23 |
| | 1.10.238.10: EF-hand | 5202 | 53 | 89 +- 45 |
| | 1.10.10.60: Homeodomain-like | 5464 | 37 | 54 +- 19 |
| | 1.10.287.470: Helix hairpin bin | 8318 | 2 | 49 +- 24 |
| **1.20: Up-Down Bundle** | 1.20.1250.20: MFS general substrate transporter-like domains | 9727 | 9 | 172 +- 95 |
| | 1.20.1070.10: Rhodopsin 7-helix | 4560 | 8 | 162 +- 83 |

| | | | | |
|---|---|---|---|---|
| | transmembrane proteins | | | |
| | 1.20.1560.10: ABC transporter type 1, transmembrane domain | 4825 | 5 | 95 +- 102 |
| **1.25: Alpha Horseshoe** | 1.25.40.10: Tetratricopeptide repeat domain | 58907 | 41 | 143 +- 68 |
| | 1.25.40.20: Ankyrin repeat-containing domain | 22379 | 14 | 136 +- 72 |
| | 1.25.10.10: Leucine-rich Repeat Variant | 14129 | 1 | 195 +- 122 |
| **2.30: Roll** | 2.30.40.10: Urease, subunit C, domain 1 | 5519 | 5 | 30 +- 19 |
| **2.40: Beta Barrel** | 2.40.50.100: RNA polymerase II/Efflux pump adaptor protein, barrel-sandwich hybrid domain | 7877 | 12 | 40 +- 21 |
| | 2.40.10.10: Trypsin-like serine proteases | 5239 | 14 | 74 +- 51 |
| | 2.40.50.140: Nucleic acid-binding proteins | 5749 | 51 | 77 +- 32 |
| **2.60: Sandwich** | 2.60.120.10: Jelly Rolls | 4235 | 24 | 99 +- 41 |
| | 2.60.120.200 | 5295 | 16 | 148 +- 56 |
| | 2.60.120.260: Galactose-binding domain-like | 6624 | 18 | 125 +- 42 |
| | 2.60.40.10: Immunoglobulins | 26258 | 94 | 92 +- 21 |
| **2.120: 6 Propeller** | 2.120.10.30: TolB, C-terminal domain | 5528 | 4 | 153 +- 56 |
| **2.130: 7 Propeller** | 2.130.10.10: YVTN repeat-like/Quinoprotein amine dehydrogenase | 25805 | 25 | 171 +- 80 |
| **2.150: 2 Solenoid** | 2.150.10.10: Serralysin-like metalloprotease, C-terminal | 6963 | 2 | 118 +- 55 |
| **2.160: 3 Solenoid** | 2.160.20.10: Single-stranded right-handed beta-helix, Pectin lyase-like | 5916 | 7 | 206 +- 96 |
| **3.20: Alpha-Beta Barrel** | 3.20.20.140: Metal-dependent hydrolases | 3803 | 18 | 80 +- 68 |
| | 3.20.20.70: Aldolase class I | 4024 | 41 | 106 +- 66 |
| | 3.20.20.80: Glycosidases | 5024 | 35 | 129 +- 89 |
| **3.30: 2-Layer Sandwich** | 3.30.200.20: Phosphorylase Kinase; domain 1 | 7455 | 38 | 48 +- 34 |
| | 3.30.450.40: GAF domain | 4217 | 10 | 133 +- 44 |

| | | | | |
|---|---|---|---|---|
| | 3.30.160.60: Classic Zinc Finger | 15979 | 37 | 38 +- 22 |
| | 3.30.450.20: PAS domain | 10945 | 13 | 100 +- 33 |
| | 3.30.420.10: Ribonuclease H-like superfamily/Ribonuclease H | 5115 | 18 | 111 +- 50 |
| | 3.30.420.40: ATPase, nucleotide binding domain | 11728 | 23 | 56 +- 44 |
| | 3.30.70.270: Reverse transcriptase/Diguanylate cyclase domain | 4066 | 7 | 68 +- 54 |
| **3.40: 3-Layer (aba) Sandwich** | 3.40.50.300: P-loop containing nucleotide triphosphate hydrolases | 27918 | 158 | 119 +- 69 |
| | 3.40.50.2000: Glycogen Phosphorylase B; | 18001 | 22 | 84 +- 64 |
| | 3.40.50.720: NAD(P)-binding Rossmann-like Domain | 19154 | 63 | 71 +- 59 |
| | 3.40.50.2300: Response regulator | 14971 | 22 | 85 +- 47 |
| | 3.40.50.1820: Alpha/Beta hydrolase fold, catalytic domain | 9428 | 33 | 131 +- 68 |
| | 3.40.1110.10: Calcium-transporting ATPase, cytoplasmic domain N | 3698 | 3 | 35 +- 40 |
| | 3.40.50.1000: HAD superfamily/HAD-like | 5364 | 19 | 61 +- 53 |
| | 3.40.30.10: Glutaredoxin | 4624 | 36 | 89 +- 42 |
| | 3.40.50.150: Vaccinia Virus protein VP39 | 8742 | 48 | 112 +- 58 |
| | 3.40.190.10: Periplasmic binding protein-like II | 21309 | 26 | 66 +- 50 |
| | 3.40.630.30: Gcn5-related N-acetyltransferase (GNAT) | 5410 | 19 | 114 +- 51 |
| **3.50: 3-Layer (bba) Sandwich** | 3.50.50.60: FAD/NAD(P)-binding domain | 8037 | 22 | 74 +- 59 |
| **3.80: Alpha-Beta Horseshoe** | 3.80.10.10: Ribonuclease Inhibitor | 23046 | 28 | 172 +- 90 |
| **3.90: Alpha-Beta Complex** | 3.90.1150.10: Aspartate Aminotransferase, domain 1 | 6004 | 20 | 44 +- 36 |

**Table S5. Description of the superfamilies in the TOP50 SUPERFAMILIES dataset**

A high-level description of the superfamilies in the TOP50 SUPERFAMILIES dataset has been mentioned in Table S5. From this table, we can infer important information such as the structural diversity of a superfamily and other details such as how conserved the sequence lengths are.

#### S2.2. ANN Optimization Study

In this section, we optimized an ANN trained on the embeddings from the ProtT5 pLM to design the CATHe model. In this study, we altered the number of layers, and the number of nodes per layer in the ANN model and measured its performance on the TOP50 SUPERFAMILIES and the TOP 1773 SUPERFAMILIES datasets. Additionally, we made inferences on how the ANN model affects its performance. Furthermore, we used this study to find the breaking point of the ANN model trained on ProtT5 embeddings. It is important to conduct this analysis to arrive at an optimal architecture for CATHe. Moreover, it serves as proof that the high performance attained by the deep learning models is not a chance occurrence and that the deep learning models are able to pick up important information in order to classify the sequences into CATH superfamilies. In Tables S6 (A), and S6 (B), we describe the 11 different ANN models (models 1-11) that were developed for the analysis, and their performances on the TOP50 SUPERFAMILIES and the TOP 1773 SUPERFAMILIES datasets, respectively.

The first five models (models 1-5) have minimal differences in their performance (Fig. S6 (B)). This shows that a 1-layer neural network with 128 nodes (model 5) is enough to reach performance similar to that of a highly computationally expensive model like model 1. Therefore, the architecture of model 5 was chosen for making CATHe. From model 6, we note a sharp decrease in performance. The poor performance of models 10 and 11 denote the breaking point of the ANN; additionally, they prove that the high performance attained by CATHe can be trusted to detect remote homologues for CATH superfamilies.

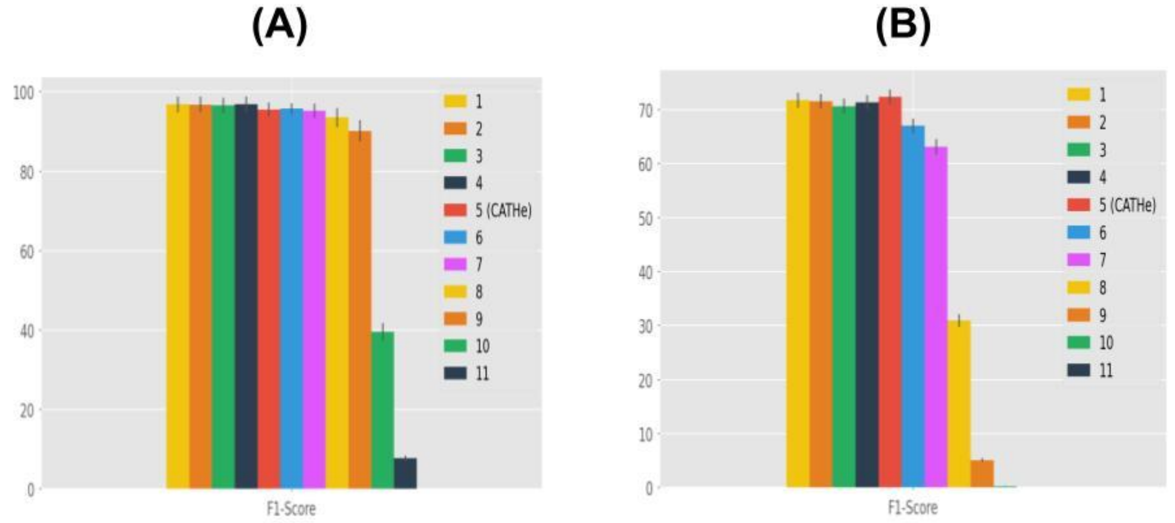

**Fig S8. ANN + ProtT5 models with descending complexity (in terms of the number of parameters, all of which have been described in Table 4) plotted against their F1-Scores on the ProtT5 embeddings of the testing set for (a) the TOP50 SUPERFAMILIES dataset, and (b) the TOP 1773 SUPERFAMILIES dataset**

| Model S.No. | Architecture | Accuracy | F1-Score | MCC | Bal Acc | Epochs | Total Parameters |
|---|---|---|---|---|---|---|---|
| 1 | 1024x2 | 98.46 +- 0.27 | 96.82 +- 1 | 0.984 +- 0.0028 | 97.07 +- 0.82 | 55 | 2,158,642 |
| 2 | 1024x1 | 98.41 +- 0.28 | 96.81 +- 1 | 0.9834 +- 0.0029 | 96.94 +- 0.82 | 53 | 1,104,946 |
| 3 | 512x1 | 98.36 +- 0.28 | 96.57 +- 1 | 0.9829 +- 0.0029 | 96.69 +- 0.85 | 48 | 552,498 |
| 4 | 256x1 | 98.4 +- 0.28 | 96.82 +- 1 | 0.9834 +- 0.0029 | 96.87 +- 0.85 | 47 | 276,274 |
| 5 | 128x1 | 98.15 +- 0.3 | 95.49 +- 0.93 | 0.9807 +- 0.0031 | 95.98 +- 0.67 | 57 | 138,162 |
| 6 | 64x1 | 97.89 +- 0.33 | 95.76 +- 0.68 | 0.9781 +- 0.0034 | 95.84 +- 0.68 | 43 | 69,106 |
| 7 | 32x1 | 97.68 +- 0.34 | 95.18 +- 0.92 | 0.9759 +- 0.0036 | 95.7 +- 0.67 | 36 | 34,578 |
| 8 | 10x1 | 97.07 +- 0.37 | 93.57 +- 1.23 | 0.9696 +- 0.0038 | 94.97 +- 0.7 | 39 | 10,840 |
| 9 | 5x1 | 94.34 +- 0.53 | 90.17 +- 1.37 | 0.9411 +- 0.0055 | 90.83 +- 1.2 | 82 | 5,445 |
| 10 | 2x1 | 54.89 +- 1.12 | 39.62 +- 1.13 | 0.5336 +- 0.0113 | 43.8 +- 1.25 | 84 | 2,208 |
| 11 | 1x1 | 25 +- 0.98 | 7.74 +- 0.36 | 0.2138 +- 0.0095 | 12.25 +- 0.58 | 44 | 1,129 |

**Table S6. (A) Description of the eleven ANN architectures, and their performances on the ProtT5 embeddings of the testing set of the TOP50 SUPERFAMILIES dataset**

| Model S.No. | Architecture | Accuracy | F1-Score | MCC | Bal Acc | Epochs | Total Parameters |
|---|---|---|---|---|---|---|---|
| 1 | 1024x2 | 85.67 +- 0.41 | 71.71 +- 0.7 | 0.8562 +- 0.0041 | 75.94 +- 0.63 | **200** | 3,924,717 |
| 2 | 1024x1 | 85.07 +- | 71.55 +- | 0.85 +- | 75.4 +- | **200** | 2,871,021 |

| | | | | | | | |
|---|---|---|---|---|---|---|---|
| | | 0.41 | 0.69 | 0.0041 | 0.63 | | |
| 3 | 512x1 | 84.98 +- 0.41 | 70.63 +- 0.7 | 0.8491 +- 0.0042 | 74.48 +- 0.63 | **200** | 1,436,397 |
| 4 | 256x1 | 85.19 +- 0.41 | 71.34 +- 0.69 | 0.8513 +- 0.0041 | 75.05 +- 0.63 | **174** | 719,085 |
| 5 | 128x1 | **85.6 +- 0.41** | **72.35 +- 0.67** | **0.8554 +- 0.0041** | **76.11 +- 0.62** | **200** | 360,429 |
| 6 | 64x1 | 83.24 +- 0.43 | 66.99 + 0.7 | 0.8317 +- 0.0043 | 71.25 +- 0.64 | **200** | 181,101 |
| 7 | 32x1 | 81.26 +- 0.45 | 63.09 +- 0.72 | 0.8118 +- 0.0046 | 67.87 +- 0.66 | **200** | 91,437 |
| 8 | 10x1 | 63.18 +- 0.56 | 30.91 +- 0.58 | 0.6302 +- 0.0056 | 36.39 +- 0.58 | **135** | 29,793 |
| 9 | 5x1 | 32.88 +- 0.57 | 5.1 +- 0.18 | 0.3249 +- 0.0057 | 7.47 +- 0.21 | **122** | 15,783 |
| 10 | 2x1 | 10.49 +- 0.36 | 0.29 +- 0.01 | 0.0964 +- 0.0034 | 0.7 +- 0.02 | **68** | 7,377 |
| 11 | 1x1 | 5.81 +- 0.27 | 0.06 +- 0.006 | 0.0491 +- 0.0025 | 0.28 +- 0.009 | **35** | 4,575 |

**Table S6. (B) Description of the eleven ANN architectures, and their performances on the ProtT5 embeddings of the testing set of the TOP 1773 SUPERFAMILIES dataset**

#### S2.3. Metrics:

A confusion matrix tabulates the actual values and the predicted values, allowing one to understand the performance of the model. The True Positive (TP), True Negative (TN), False Positive (FP), and False Negative (FN) parameters of the confusion matrix were used to calculate all performance metrics. TP refers to those data points for which the model correctly predicts the proper positive class, and FP refers to those data points for which the model falsely predicts the positive class. In that same sense, TN refers to those data points for which the model correctly predicts the proper negative class, and FN refers to those data points where the model falsely predicts the negative class. The four metrics have been delineated below.

##### a) Accuracy

Accuracy is defined using the formula below:

$$Accuracy = (TP + TN) / (TP + FP + TN + FN)$$

This accuracy is calculated for each of the classes individually and then the average of these values is taken as the accuracy of the model. This metric is frequently used for measuring model performance, but there is a caveat. This metric is quite sensitive to class imbalance, and as the datasets we are using in this study are significantly imbalanced, this metric would not be able to display the true performance of the model.

###### b) F1-Score

F1-Score is the harmonic mean of precision and recall; the formulae have been outlined below:

$$Precision = TP / (TP + FP)$$
$$Recall = TP / (TP + FN)$$
$$F1 - Score = (2 * Precision * Recall) / (Precision + Recall)$$

This F1-Score is calculated for each of the classes individually and then the average of these is taken as the F1-Score of the model.

###### c) Matthews Correlation Coefficient (MCC)

MCC is one of the most robust metrics that can be used to measure the performance of a classification model, especially when the dataset used is affected by class imbalance. The formulae for calculating this metric have been outlined below:

$$N = TP + FP + FN + TN$$
$$S = (TP + FN) / N$$
$$P = (TP + FP) / N$$
$$MCC = (TP/N - S * P) / \sqrt{PS(1 - S)(1 - P)}$$

This MCC is calculated for each of the classes individually and then the average of these is taken as the MCC for the model.

###### d) Balanced Accuracy (Bal Acc)

Balanced accuracy for this multi-class classification scenario is defined as the average of the recall for each individual class.

#### S2.4. SSAP Thresholds for Structural Validation

To determine which thresholds to use when we structurally compare domains to their structural representatives in the CATHe predicted superfamilies, we conducted a threshold analysis using SSAP. For this analysis, we used domains from the Golden Benchmark dataset. This dataset consisted of domains that had the same annotations in CATH and SCOP. This dataset was clustered at 30% sequence identity and the resultant set had 3,186 domains. We conducted an all-vs-all structure comparison on these S30 representatives of the Golden Benchmark dataset. As we know which domain pairs were homologues and which were not, we used that information to plot the SSAP scores at a fixed SSAP overlap (60%) for the different CATH classes (1, 2, and 3) at different error rates (Figure S6). From the plot, we extracted SSAP score thresholds at a 5% error rate for the different CATH classes, and they are as follows:

- Class 1: SSAP score of 71, and SSAP overlap of 60%
- Class 2: SSAP score of 66, and SSAP overlap of 60%
- Class 3: SSAP score of 69, and SSAP overlap of 60%

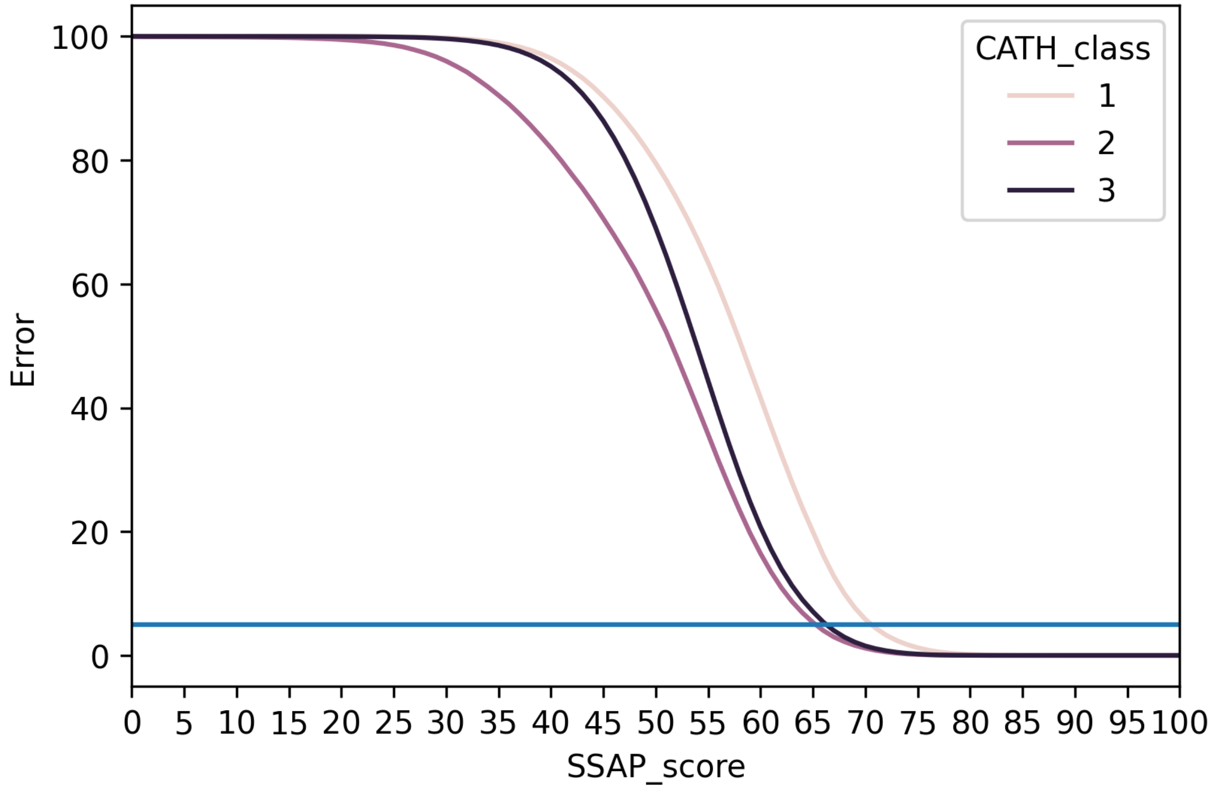


**Fig S9. SSAP score values were plotted against error rates for three different CATH classes, 1 (Mainly Alpha), 2 (Mainly Beta), and 3 (Alpha Beta). The blue line shows the mark for a 5% error rate.**

#### S2.5. Models

| **Model Name** | **Brief Description** |
|---|---|
| BLAST | Protein sequence homology based inference using BLAST |

| | |
|---|---|
| ANN + Length | Artificial Neural Network model trained on protein sequence lengths |
| Random | Random prediction model with random superfamily annotation |
| CATHe | Artificial Neural Network model trained on embeddings from ProtT5 pLM |
| ANN + ProtBERT | Artificial Neural Network model trained on embeddings from ProtBERT pLM |
| LR + ProtBERT | Logistic Regression model trained on embeddings from ProtBERT pLM |
| LR + ProtT5 | Logistic Regression model trained on embeddings from ProtT5 pLM |

**Table S7. Description of the seven different prediction models that were developed in this study to detect remote homologues for CATH superfamilies**
